# Zinc coordination by the wheat rust effector AvrSr33 provides a structural scaffold required for immune recognition

**DOI:** 10.64898/2026.06.24.731497

**Authors:** Zhao Li, Jian Chen, Emily M. Cross, Megan A. Outram, Daniel S. Yu, Jana Sperschneider, Melania Figueroa, Peter N. Dodds, Simon J. Williams

## Abstract

Interactions between plants and pathogens drive long-term co-evolution through cycles of effector diversification and immune recognition. Effectors play central roles in this molecular interplay, and zinc-binding folds have been identified in a subset of pathogen effectors, yet the contribution of zinc coordination to effector stability and immune recognition remains unclear.

Here, we investigated the structure and recognition of the AvrSr33 effector from the wheat stem rust fungus (*Puccinia graminis* f. sp. *tritici*, *Pgt*). Structural and biochemical analyses show that AvrSr33 adopts a fold containing two zinc-binding sites.

Transient expression assays in *Nicotiana benthamiana* show that AvrSr33 directly interacts with Sr33, and comparison of recognised and non-recognised AvrSr33 variants identifies a polymorphic loop associated with recognition. This loop is positioned adjacent to one of the zinc-binding sites with its orientation constrained by zinc coordination. Reciprocal mutations of key surface residues within this region alter recognition, whereas mutation of the zinc-binding site abolishes recognition.

Our data suggest that zinc coordination in AvrSr33 provides a structurally constrained scaffold that supports the surface features associated with Sr33 recognition. These findings provide a mechanistic framework for understanding how zinc coordination contributes to effector recognition and may influence the evolutionary trajectories of pathogen effectors.

## Introduction

Global food security is under constant threat from plant diseases. To infect plants, pathogens deliver an array of secreted proteins, termed effectors, into the apoplast or directly inside host cells, where they manipulate host physiology and suppress immune responses (Jones and Dangl 2006; Lo Presti et al. 2015). In response, plants have evolved resistance genes encoding immune receptors that specifically detect effectors and trigger a strong immune response, known as effector-triggered immunity (ETI) (Ngou, Ding, and Jones 2022; Dodds, Chen, and Outram 2024). Recognised effectors are referred to as avirulence proteins (Avr). Recognition imposes strong selective pressure on pathogens, driving rapid effector diversification, which in turn poses a major challenge for disease management. Therefore, characterising the molecular features that govern effector recognition and evasion is critical for understanding how plant–pathogen interactions shape the evolution of immune receptor specificity.

Wheat stem rust, caused by the fungus *Puccinia graminis f. sp. tritici* (*Pgt*), is among the most devastating wheat diseases, particularly since the emergence of the highly virulent *Pgt* lineage Ug99 (Pretorius et al. 2000; Hovmoller, Walter, and Justesen 2010). Like other rust fungi, *Pgt* is a biotrophic pathogen that delivers effectors into host cells via specialised infection structures called haustoria (Henningsen et al. 2025). Multiple stem rust resistance (*Sr*) genes have been identified in wheat (Dracatos et al. 2023), most of which encode canonical coiled-coil nucleotide-binding leucine-rich repeat proteins (CNLs), a major class of nucleotide-binding leucine-rich repeat (NLR) receptors. To date, seven Avrs that are recognised by stem rust CNLs have been identified (Arndell et al. 2024; Chen et al. 2017; Salcedo et al. 2017; Upadhyaya et al. 2021; Chen et al. 2025; Spanner et al. 2026). Of these, the structure of AvrSr35, ArvSr50 and AvrSr27 effectors have been determined experimentally, supporting studies that have characterised their recognition by corresponding CNL receptors. The stem rust effector AvrSr35 directly binds and activates the Sr35 receptor, which assembles into a pentameric resistosome (Forderer et al. 2022; Zhao et al. 2022). The interaction interface is localised to the α10 helix of AvrSr35 and the ascending lateral side of the last eight LRR units of Sr35. AvrSr50 is also directly recognised by Sr50, and a single amino acid substitution within the effector is sufficient for the protein to evade recognition (Ortiz et al. 2022). AvrSr27 is composed of two duplicate zinc-bound domains, whereby its N-terminal domain alone is sufficient for recognition and association with Sr27 (Outram et al. 2024). In contrast to these direct NLR recognition systems, recent work revealed that the Sr62 resistance locus encodes a tandem kinase-NLR module that functions as a two-component immune receptor. The effector AvrSr62 binds to the N-terminal domain of the tandem kinase, disrupting intramolecular interactions within this protein and thereby activating the associated NLR (Chen et al. 2025).

Fungal effectors are often rich in cysteines, and their high frequency is used as a distinguishing feature (Sperschneider et al. 2026). Given the contrasting redox environments of the apoplast and cytoplasm, cysteine-rich effectors may employ different structural strategies. In the oxidizing apoplast, cysteines within fungal effectors have been shown to stabilise protein structure via the formation of disulfide bonds (Doehlemann and Hemetsberger 2013; Outram et al. 2021). In a reducing cytoplasmic environment, their structure could be stabilized through metal ion coordination (Jennings et al. 2018). Structure determination of three cysteine-rich cytoplasmic fungal effectors, AvrPii from the rice blast pathogen *Magnaporthe oryzae* (De la Concepcion et al. 2022), AvrP from the flax rust pathogen *Melampsora lini* (Zhang et al. 2018), and AvrSr27 from the stem rust pathogen *Pgt* (Outram et al. 2024), have shown zinc-binding with cysteine-mediated coordination. Recently, we also demonstrated biochemically that two additional recognised effector proteins from Pgt, AvrSr13 and AvrSr22, bind zinc (Outram et al. 2026). Structural predictions of these also suggest the metal binding is coordinated by cysteine residues. In the case of AvrP, zinc ion binding was required for recognition by its cognate flax resistance protein, P (Zhang et al. 2018) but it remains unclear how zinc binding contributes to recognition or interaction with receptors or host targets. Another Pgt effector, AvrSr33, was recently identified (Spanner et al. 2026) and is recognised by the wheat Sr33 protein (Periyannan et al. 2013), although the molecular basis of recognition remains unclear. AvrSr33 is cysteine-rich and it is possible that the cysteine residues may also contribute to structural stability through zinc ion coordination. Eight variants of AvrSr33 have been identified in *Pgt* genomes (Spanner et al. 2026). AvrSr33-01 to 04 are recognised by Sr33, while AvrSr33-05 to 08 are not.

In this study, using biochemical and structural analyses we demonstrate that AvrSr33 is a zinc-binding effector that coordinates two zinc ions via cysteine residues. *In planta* interaction assays suggest that AvrSr33 directly interacts with Sr33. We identified the surface region of AvrSr33 that is required for interaction with Sr33 through site-directed mutagenesis, guided by comparisons to non-recognised AvrSr33 variants. This region co-localises with one of the zinc binding sites. These findings reveal that zinc coordination can shape effector recognition in plant-pathogen interactions.

## Materials and Methods

### Cloning

For recombinant protein production, the cDNA of AvrSr33-01 and AvrSr33-05 lacking the signal sequence predicted by SIGNALP-6.0 (Teufel et al. 2022), corresponding to amino acids 21-124, were used. The sequences were codon-optimised for *Escherichia coli* expression and double-stranded DNA fragments were synthesised by Integrated DNA Technologies Inc. (IDT®, Coralville, IA, USA). DNA fragments were cloned into a modified pOPIN vector (Bentham AR 2021) using Golden Gate cloning, resulting in a construct encoding an N-terminal SUMO-6His tag followed by a Human Rhinovirus (HRV) 3C protease cleavage site to allow 6His tag removal. All constructs were cloned with a 3’ stop codon after the coding sequence. Protein sequences following purification and tag-cleavage are listed in Supporting Information Table S1.

For transient expression of AvrSr33 and mutant variants in *Nicotiana benthamiana*, the effector constructs were initially cloned into the pDONR^TM^207 entry vectors by BP reactions (Invitrogen) and then into the pBIN19-35S-YFP-GWY destination vectors by LR reactions (Invitrogen) (Earley et al. 2006). Multiple mutagenesis fragments of AvrSr33 were obtained from IDT. Site-directed mutagenesis was conducted using the Phusion High-Fidelity DNA Polymerase (M0530L; NEB), following by DpnI treatment to erase the methylated plasmid template. Constructs and primer sequences are detailed in Table S2. Sr33 was cloned into the pTK009-pUbi-YFP vector (Ortiz et al. 2022). For the split-GAL4 RUBY assay, Sr33 was inserted into the pAM-35S-GAL4BD-GWY vector, the α1 helix of Sr33 was removed to eliminate cell death executed by Sr33. The AvrSr33 variants were cloned into the pAM-35S-VP16-GWY vector using Gateway cloning (Chen et al. 2023). Plasmid integrity was confirmed using Sanger sequencing. Multiple sequence alignments were generated using Clustal Omega and visualized with ESPript v3.0 (Robert and Gouet 2014).

### Protein production

Proteins were expressed in SHuffle T7 Express lysY Competent *E. coli* (C3030J) or in BL21 T7 Express lysY competent *E. coli* (C3010I; NEB). Bacterial cultures were grown in Terrific Broth (TB) medium supplemented with trace elements (1000x trace elements solution containing 50 mM FeCl_3_, 20 mM CaCl_2_, 10 mM MnCl_2_, 10 mM ZnSO_4_, 2 mM CoCl_2_, 2 mM CuCl_2_, and 2 mM NiCI_2_) (Studier 2005) at 37°C with shaking at 220 RPM until they reached to OD600 = 0.6–0.8, after which they were induced with the addition of 200 μM isopropyl-1-thio-b-D-galactopyranoside (IPTG). After induction, cultures were incubated for a further 16 h at 16°C and then harvested by centrifugation at 3500 x *g* for 15 min at 4°C. Cells were resuspended in a lysis buffer containing 50 mM HEPES, 500 mM NaCl, 10% glycerol, 1 mM phenylmethanesulfonylfluoride (PMSF). For the BL21 grown cells, 1 mM dithiothreitol (DTT) was supplemented into the lysis buffer. Cells were lysed on ice by sonication (5 min, 40% amplitude, 10 s/on, 10 s/off), followed by centrifugation at 20,000 x *g* for 40 min at 4°C. The clarified lysate was applied to a 5 mL HisTrap FF column (Cytiva, Uppsal, Sweden) and then washed with 20 column volumes (c/v) of wash buffer (50 mM HEPES, 300 mM NaCl, and 30 mM imidazole). The proteins were eluted using a linear gradient with over 10 c/v elution buffer (50 mM HEPES, 300 mM NaCl, 300 mM imidazole). The fractions were visualised by Coomassie-stained SDS-PAGE and those containing the target protein were collected into a 3.5 kDa SnakeSkin^TM^ dialysis tube (Thermo Fisher Scientific, Waltham, MA, USA). HRV 3C protease was added to the collected protein solution before putting into the dialysis buffer (10 mM HEPES with 300 mM NaCl) at 4°C overnight. The removed tag sample was purified further by size exclusion chromatography (SEC, Superdex 75 increase 10/300GL column, Cytiva) with SEC buffer (10 mM HEPES with 150 mM NaCl). All buffers used for purification of AvrSr33-01 were adjusted to pH 7.5, whereas buffers for AvrSr33-05 were adjusted to pH 7.2. Proteins were concentrated using a 3 kDa molecular weight cut-off Amicon centrifugal concentrator (MilliporeSigma, Massachusetts, USA). The purity of the protein samples was analysed using SDS-PAGE and protein concentrations were determined using a SpectraMax QuickDrop UV-Vis Spectrophotometer (Molecular Devices, San Jose, CA, USA) with absorbance at 280 nm. The extinction coefficient used for protein concentration determination was calculated using the Expasy ProtParam tool: http://ca.expasy.org/cgibin/protparam/.

### Intact mass spectrometry

The monoisotopic mass of AvrSr33-01 and AvrSr33-05 in both native and reduced states were detected by an Orbitrap Fusion™ Tribrid™ mass spectrometer (Thermo Fisher Scientific, Massachusetts, USA) coupled with Agilent UHPLC system (Agilent, California, USA). All samples were adjusted to a final concentration of 10 mM in 0.1% (v/v) formic acid (FA) for detection. Native samples were analysed directly, while for reduced samples, 10 mM DTT was added to the protein solution followed by incubation for 30 min at 60°C. For each sample, 5 μL was loaded for measurement. Each sample was first desalted for 2 min on an Agilent C3 trap column (ZORBAX StableBond C3) at a flow rate of 500 µL/min at 95% buffer A (0.1% v/v FA) and 5% buffer B (0.1% v/v FA and 100% acetonitrile) followed by separation over 8 min using a 5–80% (v/v) gradient of buffer B at a flow rate of 500 µL/min. The eluted protein samples were ionized by heated electrospray ionisation to form charged species and introduced into the mass spectrometer for m/z analysis. The acquisition was performed across m/z 200-4000 with an accumulation time of 1 s. Freestyle software (Thermo Fisher Scientific) was used to analyse the results. Theoretical monoisotopic masses were calculated using the Expasy PeptideMass program (https://web.expasy.org/peptide_mass/).

### 4-(2-pyridylazo) resorcinol (PAR) assay

PAR assays were performed as described previously (Zhang et al. 2017). The PAR solution was prepared in MilliQ water to a concentration of 1 mM. Protein samples (10 μM) were boiled with 1% (w/v) SDS for 5 min at 95°C for metal ion release and then PAR added to a final concentration of 50 μM. The SEC buffer was used as a blank measurement. The non-metal binding effector AvrSr50 expressed from *E. coli* (Ortiz et al. 2022) was used as a negative control. Samples absorbance was measured using a Spectromax Quickdrop (Molecular Devices, San Jose, CA, USA) over a wavelength range of 300-700nm.

### Inductively coupled plasma spectrometry (ICP-MS)

ICP-MS was used to quantify the metal ions in the protein samples, following the method described previously (Outram et al. 2024; Martinez-D’Alto et al. 2023). Protein samples (5 μM) were treated with 2% (v/v) nitric acid (HNO_3_) at room temperature overnight. The samples were then centrifuged, and the supernatant was diluted ten-fold with 2% (v/v) HNO_3_ before analysis by ICP-MS. Agilent Intelliquant 68 multi-element standard No. 1 (IQ-1, #5190-9422) at 100 µg mL^-1^ in 5% HNO_3_ was used to prepare standards for the calibration curve. Protein samples were analysed using a Thermo Fisher iCap RQ Quadrupole (Thermo Fisher Scientific) ICP-MS in Kinetic Energy Dispersion Sensitive mode with helium as a collision gas at 1550 W. Argon was used for plasma generation and sample nebulisation. Data were processed using Qtegra^TM^ software. Limit of detection (LOD) was calculated from the standard deviation (SD) across instrument blanks in triplicates. Each experiment was carried out in triplicate.

### Crystallization and structure determination

For determining the crystallisation conditions of AvrSr33-01, initial screening was performed in MRC2 96-wells plates at 20°C using the sitting drop vapour-diffusion method and commercially available sparse matrix screens. The drops, consisting of 200 nL protein solution and 200 nL reservoir solution, were equilibrated against 40 μL reservoir solution. All dispensing steps were performed using an NT8®-Drop Setter robot (Formulatrix, Bedford, MA, USA), and the drops were monitored and imaged using the Rock Imager system (Formulatrix). Crystal hits were obtained in several screen conditions. The best crystals were further optimised by grid screening using 24-well hanging-drop vapour-diffusion plates, with the hanging drop consisting of 1 µL protein solution and 1 µL of reservoir solution, over 500 µL reservoirs. The final condition used to obtain diffraction quality crystals for AvrSr33-01 (5 mg/mL) consisted of 0.2 M magnesium acetate tetrahydrate and 20% (w/v) PEG 3350, with the protein in 10mM HEPES (pH 7.5). The crystals were transferred into a cryoprotectant solution containing the reservoir solution supplemented with 10% (v/v) glycerol and 10% (v/v) ethylene glycol before flash-cooling in liquid nitrogen.

Diffraction data was collected at the high performance macromolecular crystallography (MX3) beamline at the Australian Synchrotron (T. T. Caradoc-Davies 2023). Initial spot finding and indexing of raw diffraction images was performed using xia2 DIALS in the CCP4 computational suite (version 9.0.008) (Beilsten-Edmands et al. 2020; Winn et al. 2011; Winter 2010; Winter et al. 2018). Following this, the data was scaled and merged using Aimless (version 0.8.2) (Evansa and Murshudova 2013). The data was cut to 1.85 Å, with a new set of R-free flags generated (5% of total reflections). Phaser MR (version 2.8.3) was used for molecular replacement using a predicted monomeric structure of AvrSr33 generated by Alphafold3 (McCoy et al. 2007; Abramson et al. 2024). The model was pre-processed using the tool available in the Phenix computational suite (version 1.21.2-5419-000) (Liebschner et al. 2019; Oeffner et al. 2022). Repeated rounds of model building in Coot (version 0.9.8.96) (Emsley et al. 2010) and refinement in phenix.refine (version 1.21.2-5419) (Afonine et al. 2012) was performed until a final model was obtained. Structure validation was carried out using MolProbity within the phenix.refine program (Davis et al. 2004). The final model coordinates and structure factors were deposited in the Protein Data Bank under the accession no 26OF. Statistics for data collection, processing, and final structure refinement are presented in Table S3. Structures were analysed using PYMOL, ESPript v3.0 (Robert and Gouet 2014) and structural comparisons were carried out using the DALI (Holm, 2020) and Foldseek (van Kempen et al. 2024) servers.

### Agrobacterium-mediated transient expression in *Nicotiana benthamiana*

*N. benthamiana* plants were grown at 25°C with a photoperiod of 16 h light and 8 h dark. Plants were used for Agrobacterium-mediated infiltrations 3-4 weeks after transplanting, as described previously (Chen et al. 2024). In brief, the constructs containing Sr33, AvrSr33 or mutant variants were transformed into the Agrobacterium GV3103_pMP 90_RK (pTK009 constructs) or GV3101_pMP90 (pBIN19 constructs) by electroporation. After cultivation and harvesting, Agrobacterium was suspended in infiltration buffer (10 mM MES pH 5.6, 10 mM MgCl_2_ and 150 μM acetosyringone) to final OD600 of 0.5 (except Ubi-Sr33 OD600=1.5-2.0). All cultures were incubated at room temperature for 2 h before infiltration into the *N. benthamiana* leaves. Cell death induction was analysed from 3 days after infiltration. Agrobacterium carrying the silencing suppressor p19 (for cell death assay) or V2 (for split-GAL4 RUBY assay) (OD600=0.1) were co-infiltrated to enhance protein expression in *N. benthamiana*. For the split-GAL4 RUBY assay, the leaves were decolorised in 100% ethanol to remove chlorophyll pigments, and betalain was subsequently extracted by immersing three pooled leaf discs (0.8 cm diameter) from three independent leaves in 0.5 mL water. The absorbance of betalain was measured at 538 nm using a Tecan Infinite® M1000Pro (Tecan) plate reader at room temperature.

### Immunoblot analysis

Protein extractions from the infiltrated leaves were separated by a 12% polyacrylamide gel using SDS-PAGE then transferred to a nitrocellulose membrane. Blots were probed with anti-GFP mouse monoclonal antibody (11814460001, Roche) for constructs with a YFP tag, then followed with a goat anti-mouse HRP conjugate (1705047, Bio-Rad). For the split-GAL4 RUBY assay, an anti-VP16 (V4388; Sigma) antibody was used as the primary antibody, followed by an anti-rabbit HRP conjugate (W4018; Promega). Protein loading was visualised by staining membranes with Ponceau S and protein signals were developed using the SuperSignal West Femto chemiluminescence kit (Pierce, Rockford, IL, USA).

## Results

### AvrSr33 is a zinc-binding effector with conserved metal coordinate sites

The eight AvrSr33 variants show an amino acid sequence identity ranging from ∼74-99% and contain a strictly conserved pattern of cysteine and histidine residues: Cx_2_-Cx_25_-Cx_7_-Cx_10_-Cx_2_-Cx_10_-Cx_3_-Hx_21_ (Fig. S1). Previously, we have shown that similar motifs found in other rust effectors facilitate metal binding (Zhang et al. 2018; Outram et al. 2024). To investigate this further, we expressed and purified AvrSr33-01 and the non-recognised variant AvrSr33-05 using *E. coli* expression systems. The proteins were expressed with an N-terminal 6xHis-Sumo solubility tag that could be removed via a HRV 3C protease cleavage site. Expression of AvrSr33-01 was performed in BL21(DE3) pLysS strain, which provides a reducing environment, and the strain SHuffle®, which supports an oxidizing environment and the formation of disulfide bonds in proteins that contain these modifications (Lobstein et al. 2012). This strategy was previously demonstrated to distinguish between metal-bound and disulfide-bonded effectors (Zhang et al. 2017). The proteins were purified to homogeneity (Fig. 1a) and subjected to intact mass spectrometry under both native and reducing conditions. The monoisotopic mass of AvrSr33-01 corresponded to the theoretical molecular weight with reduced cysteine sidechains, regardless of the expression strain used (Fig. S2). These results indicate that AvrSr33-01 protein is present in fully reduced forms in both expression strains and do not contain disulfide bonds. The monoisotopic mass of AvrSr33-05 also demonstrated the protein was produced in a form with reduced thiol groups (Fig. S2)

**Figure 1:**
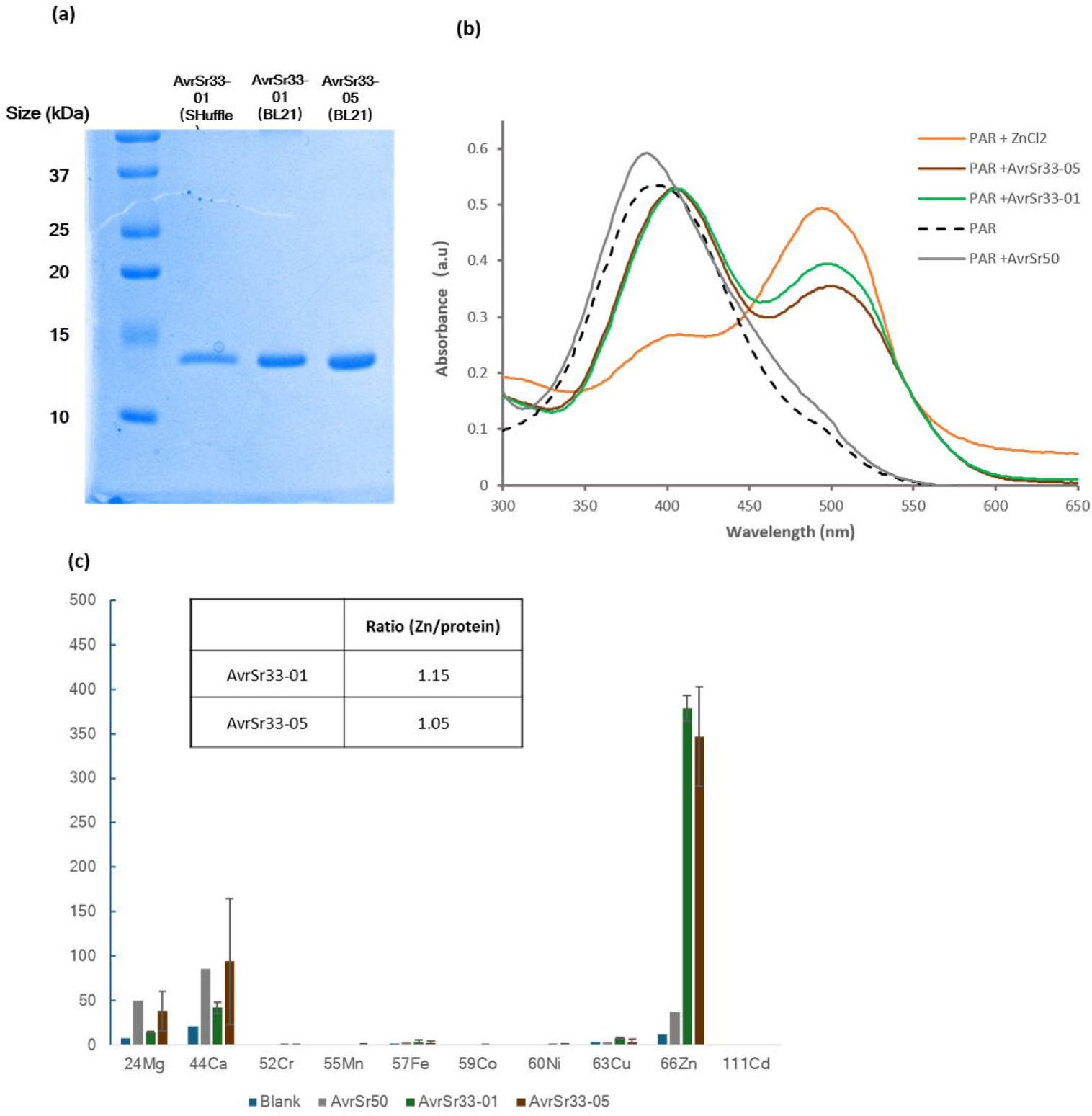
AvrSr33 is a metal-bound effector with a preference for zinc: (a) Coomassie-stained SDS–PAGE gel analysis of purified AvrSr33 proteins expressed in the *E. coli* (AvrSr33-01 in SHuffle and BL21, AvrSr33-05 BL21(DE3) pLysS. (b) Detection of metals in AvrSr33 using the 4-(2-pyridylazo) resorcinol (PAR). The absorbance spectra (300–650 nm) are shown for the following samples: PAR only, PAR plus AvrSr50 (negative control), PAR plus ZnCl2 (20 µM, positive control), PAR plus AvrSr33-01 and PAR plus AvrSr33-05. Protein produced for this assay were expressed using BL21(DE3) pLysS E. coli cells. (c) Quantification of metal ions (including Mg, Ca, Cr, Mn, Fe, Co, Ni, Cu, Zn, and Cd) in protein samples using ICP-MS. Protein samples (5DμM) were digested overnight at room temperature in 2% (v/v) HNOD prior to analysis. “Blank” indicates the buffer used for protein purification. Error bars represent the standard deviation (SD) across three replicate injections. An inset table summarizes the calculated zinc-to-protein molar ratios, offering a quantitative measure of zinc binding per protein molecule.

To investigate if the purified proteins contained bound metal ions, we used the chromogenic chelator 4-(2-pyridylazo) resorcinol (PAR) assay to detect PAR-metal complexes. A maximum absorption wavelength shift from 416 nm to 500 nm is observed when PAR forms a complex with zinc (Sabel, Shepherd, and Siemann 2009; Outram et al. 2024) (Fig. 1b). Similarly, a shift in the absorbance peak of PAR was also observed when added to denatured AvrSr33-01 or AvrSr33-05 protein solutions, indicating the release of metal ions from those proteins. Importantly, no metal ion signature was associated with AvrSr50, a non-metal binding effector protein (Ortiz et al. 2022) when purified under the same condition. To resolve the identity of the bound metal ions and estimate the stoichiometry in AvrSr33-01 and AvrSr33-05, we conducted inductively coupled plasma-mass spectrometry (ICP-MS) on the purified protein samples (Wilschefski and Baxter 2019). This analysis demonstrated a selectivity for zinc and suggested an average occupancy of ∼1 zinc ion per protomer of AvrSr33-01 and AvrSr33-05 (Fig. 1c).

### Crystal structure of AvrSr33 with zinc bound

To understand the structural basis of the zinc coordination and the overall structure of AvrSr33, we determined the crystal structure of AvrSr33-01 at 1.85 Å resolution. The Matthews coefficient was calculated and suggested 1 molecule in the asymmetric unit, with 41.26 % total solvent content. Following molecular replacement, model building and refinement, the final structure yielded an R_work_ and R_free_ of 23.3% and 26.8%, respectively (Table S3).

The structure of AvrSr33-01 consists of a compact fold centred around a four stranded antiparallel β-sheet (β1-β4), with zinc-coordinating regions positioned at either end of the β-sheet, giving the protein an overall dumbbell-like appearance (Fig. 2a). One zinc-coordinated region is formed by residues contributed from the β2-β3 turn and the N-terminal helical region, whereas extended linker regions connecting β1 to β2 and β3 to β4 form the second zinc-binding site. The structure contains two prominent α-helices (α1 and α2) within the N-terminal region, as well as a short helical turn (α3) associated with the second zinc-coordinated region. Flexible linker regions are present between α1 and α2 as well as just prior to the first β-strand. Due to intrinsic flexibility in this region, amino acids 32-33 could not be placed due to poor observed electron density.

**Figure 2:**
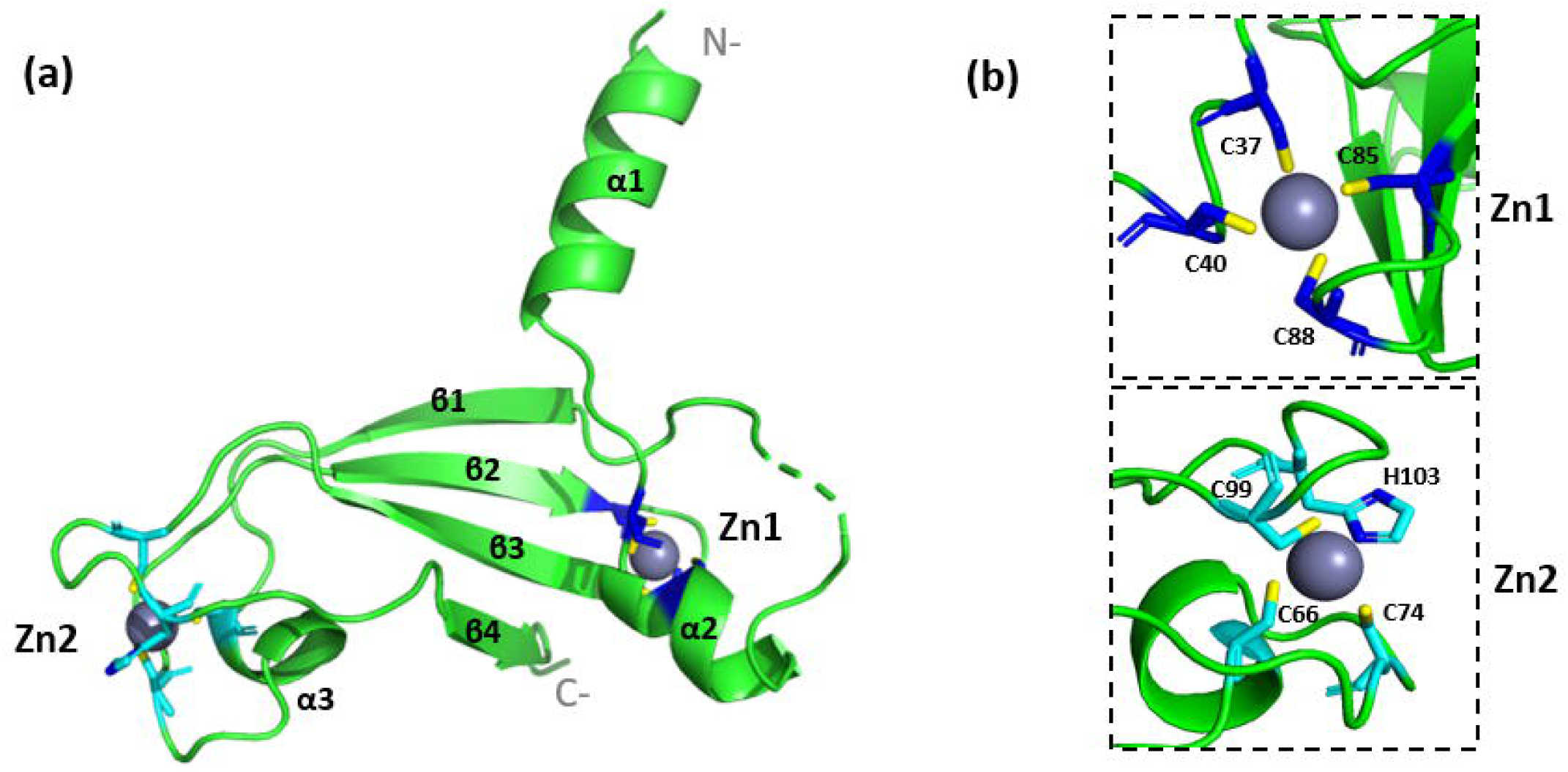
The zinc bound crystal structure of AvrSr33: (a) Cartoon representation of the overall structure of AvrSr33-01. The two bound Zn ions are shown as grey spheres with coordinating residues displayed as sticks and colored dark blue for the Zn1 site and cyan for Zn2 site. (b) The structure details of tetrahedral coordination of Zn ions. Zn1 is coordinated by a CCCC residue set, whereas Zn2 is coordinated by a CCCH residue set.

The structure revealed two metal coordination sites that are modelled with zinc ions, in agreement with biochemical assays performed (Fig. 2b). Each zinc ion is coordinated in a tetrahedral geometry, either by four cysteine residues (CCCC), we define as the zinc 1 binding site (Zn1), or by three cysteines and one histidine (CCCH), we define as the zinc 2 binding site (Zn2) (Fig 2b).

### Zinc-binding sites of AvrSr33 contribute to recognition by Sr33

We previously demonstrated that mutation of the zinc binding sites in the flax rust effector AvrP abolishes recognition by the corresponding resistance protein, P (Zhang et al. 2018). To test whether the integrity of the zinc-binding sites of AvrSr33-01 are required for recognition by Sr33, we designed single and double cysteine mutations in both zinc-binding sites. These proteins expressed were transiently co-expressed with Sr33 in *N. benthamiana*. Co-expression of AvrSr33-01 wild type (WT) protein with Sr33 resulted in a cell death response consistent with recognition of the effector by the resistance protein (Fig. 3, S3). Infiltration assays demonstrated that mutations in the Zn1-binding site of AvrSr33-01 led to a reduction in cell death intensity. The single mutation C40S resulted in weaker cell death compared with the AvrSr33-01 WT. The double mutations (C37S/C40S and C40S/C85S) abolished cell death (Fig. 3a, left). Western blot analysis demonstrated that the single mutant and the C37S/C40S double mutant accumulated in plant cells; however, the C40S/C85S mutant accumulated at much lower levels than the AvrSr33-01 WT (Fig. 3b). We subsequently investigated the impact of mutations on the Zn2-binding site. Infiltration assays showed that both single (C66S) and double (C66S/C74S) mutations completely abolished Sr33-mediated recognition (Fig. 3a, right). These visible phenotypes were consistent with quantitative cell death scores, which showed a stepwise reduction from the wild type to single and double Zn1-binding site mutations (Fig. 3c). In contrast, Zn2-site mutations scored close to zero, confirming complete loss of recognition by Sr33 (Fig. 3c). These data show that mutation of either zinc-binding site compromises Sr33 mediated recognition.

**Figure 3:**
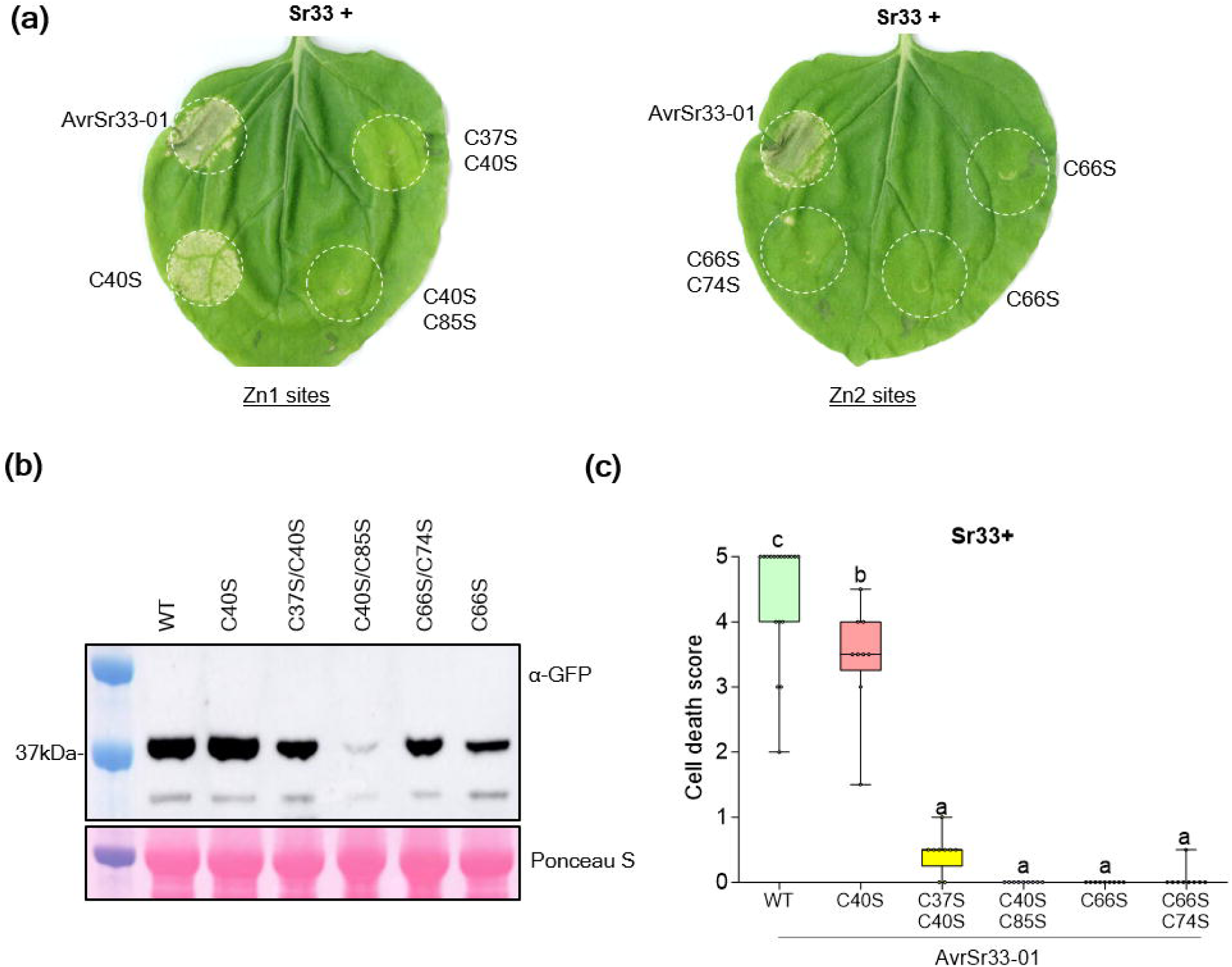
Zinc-binding sites of AvrSr33 contribute to recognition by Sr33: (a) The resistance protein Sr33 was co-expressed via agroinfiltration in *N. benthamiana* leaves with YFP-AvrSr33-01 wildtype (WT) or its cysteine to serine mutants targeting the two zinc-binding sites. Mutations include C40S, C37S/C40S and C40S/C85S in the Zn1-binding site (left leaf), and C66S and C66S/C74S in the Zn2-binding site (right leaf). White circles indicate infiltration sites. (b) Immunoblots of AvrSr33-01 WT and mutant proteins expressed in *N. benthamiana* and detected with an anti-GFP antibody. Ponceau S staining Rubisco large subunit (rbcL) was used as a loading control. (c) Boxplot graphs of cell death intensity for each treatment in (a) scored on a scale of 0–5 from at least 9 independent infiltration sites. Boxes indicate the interquartile range, the horizontal line represents the median, and whiskers show the full range of data. Different letters (a, b, c) indicate statistically significant differences among constructs (one-way ANOVA with Tukey’s HSD, p < 0.05).

### The Zn2-proximal region mediates the direct interaction between Sr33 and AvrSr33

The non-recognised AvrSr33-05 protein contains 21 amino acid polymorphisms relative to AvrSr33-01 (Fig. 4a). When these polymorphic amino acid residues were mapped on the AvrSr33-01 structure, they were found to be spread over the surface of the protein. We divided these into three spatially defined groups based on their location in either the central β-sheet (β-sheet; residues 31, 57-58, 61, 82, 93-94, 120) or the Zn1-proximal (ZP1; residues 38, 41, 50, 52, 87) or Zn2-proximal (ZP2; residues 63-65, 70, 72, 77, 97-98) regions (Fig. 4b). Based on these groups, we swapped polymorphic residues between AvrSr33-01 and AvrSr33-05 and tested their recognition by Sr33 using agroinfiltration in *N. benthamiana.* Substitution of the ZP2 residues within AvrSr33-01 to the residues found in AvrSr33-05 (AvrSr33-01 ZP2) abolished recognition by Sr33, whereas the reciprocal substitutions in AvrSr33-05 largely restored recognition of this protein (Fig. 4c left). The quantitative score analysis shown the ZP2 mutations of AvrSr33-05 was significantly higher than the non-recognised variants (P<0.0001) yet remained slightly lower than AvrSr33-01 (P<0.01) (Fig. 4c right, Fig. S4). By contrast, substitution of the polymorphic amino acids in the β-sheet (eight residues) or the ZP1 region (five residues) showed recognition patterns indistinguishable from the respective wild type AvrSr33-01 or -05 effectors (Fig. 4c). Immunoblotting showed that all the AvrSr33 protein variants were expressed, although the ZP1 region mutant of AvrSr33-01 showed lower accumulation (Fig. S4d).

**Fig. 4.**
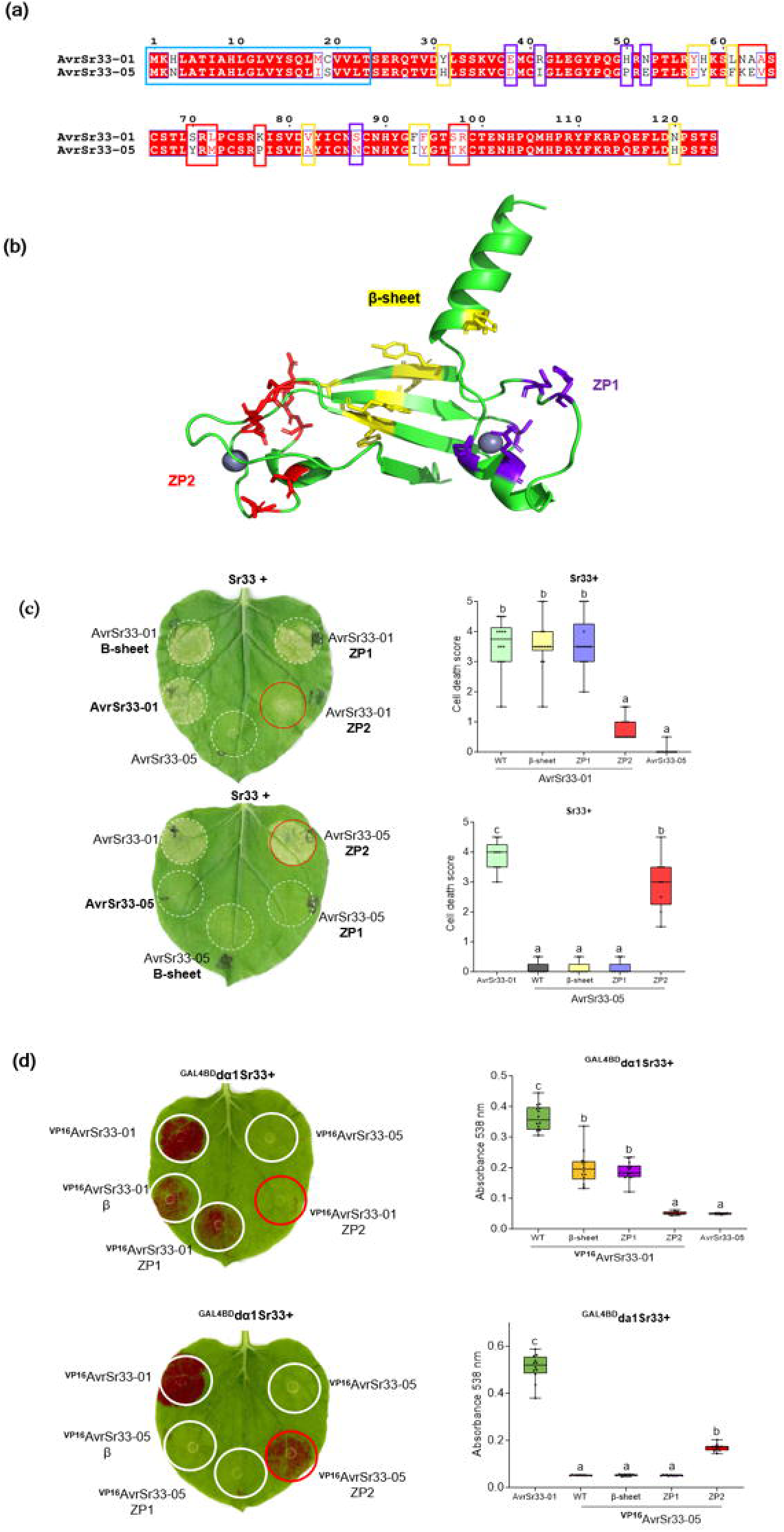
The Zn2-proximal region mediates Sr33–AvrSr33 recognition: (a) Sequence alignment of AvrSr33-01 and AvrSr33-05 illustrated by ESPript (Gouet, Robert et al. 2003). Conserved residues are shaded in red. The blue box marks the single peptide region. The yellow, orange and red box highlight the polymorphic residues located in the β sheet, Zn1-proximal (ZP1) and Zn2-proximal (ZP2) regions, respectively that were substituted between AvrSr33-01 and AvrSr33-05 variants.(b) The structure of AvrSr33-01 is shown in cartoon format with polymorphic residues from the sequence alignment indicated as sticks and coloured according to their locations: residues in the ZP1 are purple, those embedded in the β-sheet are yellow and residues in the ZP2 are red. (c) *N. benthamiana* leaves were infiltrated with Agrobacterium carrying AvrSr33-01, AvrSr33-05, or their variants with mutations in the ZP1, β-sheet, or ZP2, co-expressed with Sr33. Left panels show representative images of infiltrated leaves 3 days post infiltration. White circles indicate infiltration sites, and red circles highlight mutations that disrupted Sr33 recognition in AvrSr33-01 or restored recognition in AvrSr33-05. Right panels show graphs of cell death intensity for AvrSr33-01 and its variants (upper) and corresponding substitutions for AvrSr33-05 (lower). Cell death intensity was scored on a scale of 0–5 from at least 9 independent infiltrations. Horizontal lines represent the median, boxes represent the interquartile range, and whiskers show the full range of data. Different letters (a, b, c) denote statistically significant differences among constructs (one-way ANOVA with Tukey’s HSD, p < 0.05). (d) *N. benthamiana* leaves co-expressing UAS-RUBY+GAL4BD-dα1Sr33 with VP16-fused AvrSr33-01, AvrSr33-05, or their variants carrying mutations in the ZP1, β-sheet or ZP2 (left). White circles indicate infiltration sites, and red circles highlight mutations that abolished recognition in the AvrSr33-01 background or restored recognition in the AvrSr33-05 background. The graph on the right shows quantification of betalain accumulation by absorbance at 538 nm. Each dot represents an individual measurement from three pooled leaf disks. Horizontal lines represent the median, boxes represent the IQR, and whiskers show the full range of data. Different letters (a, b, c) denote statistically significant differences among constructs (one-way ANOVA with Tukey’s HSD, p < 0.05).

To further validate these recognition patterns at the protein-protein interaction level, we used the split-GAL4 RUBY reporter system (Chen et al. 2023). Sr33 lacking the α1 helix (da1-Sr33, 1-26 aa deletion) was fused to the GAL4 DNA binding domain (GAL4BD) and the AvrSr33 effector variants were fused to the VP16 transcription activation domain and co-expressed with the UAS-Ruby construct in *N. benthamiana.* Co-expression of VP16-AvrSr33-01 with GAL4BD-da1-Sr33 triggered strong accumulation of the purple pigment betalain (Fig. 4d), indicating physical interaction of these two proteins. The ZP2 substitutions in AvrSr33-01 completely abolished interaction, while the β-sheet and ZP1 substitutions retained betalain production but were reduced compared to the AvrSr33-01 WT. The reciprocal ZP2 substitutions in AvrSr33-05 partially activated UAS-Ruby expression, while the β-sheet and ZP1 substitutions had no effect (Fig. 4d left). Quantitative absorbance measurements at 538 nm supported these observations: signals from the ZP2 substitution mutants of AvrSr33-05 were significantly higher than those of non-recognised variants yet remained significantly below signal for the wild type (Fig. 4d right). Protein accumulation of all AvrSr33 variants were confirmed by immunoblotting (Fig. S5). Together, the cell death and split-GAL4 RUBY assay indicate that amino acids in the ZP2 region of AvrSr33-01 are required for Sr33 recognition.

### Reciprocal residue substitutions determine avirulence and virulence of AvrSr33

Previous studies have shown that a single amino acid substitution in AvrSr50 is sufficient to abolish recognition by Sr50 (Ortiz et al. 2022). To identify which of the eight polymorphic residues within the ZP2 region of AvrSr33-01 is critical for Sr33 recognition, single and combined reciprocal substitutions were introduced at this region. We further subdivided the ZP2 region into four groups and introduced reciprocal substitutions into AvrSr33-01 (NAA^63-65^/KEV, SL^70,72^/YM, K^77^/P, SR^97,98^/TK) and AvrSr33-05 (KEV/NAA, YM/SL, P/K, TK/SR) (Fig. 5, Fig. S6a&b). Substitution of SL/YM of AvrSr33-01 abolished recognition by Sr33, whereas the reciprocal replacement of YM/SL in AvrSr33-05 restored recognition by Sr33 (Fig. 5b). To confirm these findings at the protein-protein interaction level, we examined the interactions between Sr33 and the key SL/YM and YM/SL variants using the split-GAL4 RUBY assay. Substitution of SL/YM in AvrSr33-01 completely abolished the interaction with Sr33, whereas the reciprocal YM/SL substitution in AvrSr33-05 partially restored the interaction with Sr33. Although the signal was weaker than observed for the wild type AvrSr33-01/Sr33 pair, quantitative measurements showed no significant difference between AvrSr33-05 YM/SL and ZP2 construct (Fig. 5c). Protein accumulation was confirmed by immunoblotting (Fig. S6d, S5). Consistent with these results, the sequence alignment of eight AvrSr33 variants demonstrates that the SL-YM polymorphism correlates with Sr33 recognition (Fig. S1). The recognised variants AvrSr33-01 to -04 contain SL, whereas the non-recognised variants AvrSr33-05 to -08 contain YM at these positions (Spanner et al. 2026). Mapping these polymorphic residues onto the AvrSr33 structure revealed that both amino acids are found on a loop adjacent to residues that coordinate the Zn2 ion. The Zn2 ion is coordinated by four residues in a CCCH configuration. The polymorphic residues are located within a short surface-exposed loop(-S^70^RL^72^P-) positioned adjacent to the Zn2-coordinated region, following a short α-helical turn that contains two Zn2-coordinating cysteine residues (Fig. S7). This suggests that this region is involved in Sr33 recognition.

**Figure 5.**
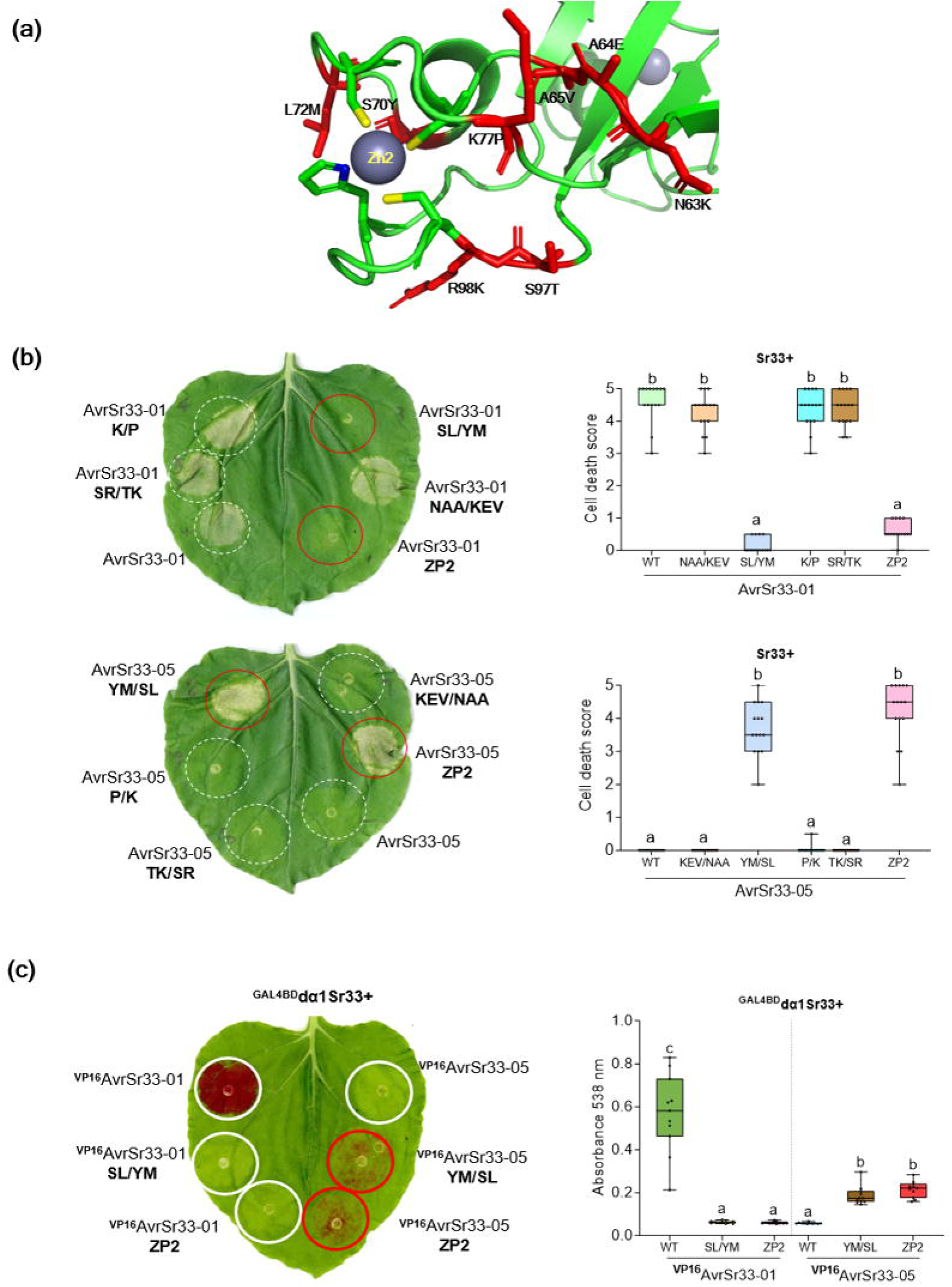
Substitution of two residues allows AvrSr33 to evade Sr33 recognition: (a) Zoomed-in view of the Zn2-proximal region of AvrSr33. Residues that differ between AvrSr33-01 and AvrSr33-05 are highlighted in red, including positions 63-65 (NAA/KEV), 70 (S/Y), 72 (L/M), 77 (K/P), and 97-98 (SR/TK) . (b) *N. benthamiana* leaves co-expressing Sr33 with AvrSr33-01, AvrSr33-05, or variants carrying mutations in the Zn2-proximal region (NAA/KEV, SL/YM, K/P, SR/TK in AvrSr33-01 and KEV/NAA, YM/SL, P/K, TK/SR in AvrSr33-05) (left). White circles indicate infiltration sites, and red circles highlight mutations that abolished recognition in the AvrSr33-01 background or restored recognition in the AvrSr33-05 background. And quantification of cell death scores (right). Graphs (right) show cell death intensity scored on a scale of 0–5 from at least n = 13 independent infiltrations. Horizontal lines represent the median, boxes represent the interquartile range, and whiskers show the full range of data. Different letters (a, b, c) denote statistically significant differences among constructs (one-way ANOVA with Tukey’s HSD, p < 0.05). (c) *N. benthamiana* leaves co-expressing UAS-RUBY+GAL4BD-Sr33 with VP16-fused AvrSr33-01, AvrSr33-05, or selected variants carrying mutations in the Zn2-proximal region (SL/YM and YM/SL respectively or the Zn2 proximal region swap) (left). White circles indicate infiltration sites, and red circles highlight functional mutations. Graph at right shows quantification of betalain accumulation by absorbance at 538 nm. Each dot represents an individual measurement from three pooled leaf disks post-infiltration. Horizontal lines represent the median, boxes represent the interquartile range, and whiskers show the full range of data. Different letters (a, b, c) denote statistically significant differences among constructs (one-way ANOVA with Tukey’s HSD, p < 0.05).

## Discussion

Dissecting how pathogen effectors are recognised or evade plant immunity offers insights into the evolutionary dynamics of pathogen effectors and provides a knowledge base for future engineering of immune receptors. Here, we explored the structural basis for recognition and evasion of the wheat stem rust effector AvrSr33-01, revealing a zinc-dependent balance between structural stability and immune activation by Sr33. Metal binding assays (PAR and ICP-MS) confirmed AvrSr33 preference for zinc binding. To assess how these sites contribute to the interaction between Sr33 and AvrSr33-01, we introduced single and double cysteine substitutions at each site and co-expressed the variants with Sr33 in *N. benthamiana*. This analysis revealed that recognition was highly sensitive to perturbation of the Zn2 site, where either single or double cysteine substitutions abolished Sr33-mediated cell death, without affecting protein accumulation in the plant. In contrast, mutations at the Zn1 site produced a graded decrease in cell death intensity, and the Zn1 double mutant (C40S/C85S) protein accumulated at markedly lower levels than the wild type. This could be explained by the structural positioning of the Zn1-coordinating residues, which act as a structural clamp that constrains local loop geometry and supports the packing of the adjacent β-strands within the protein scaffold. These results suggest that the Zn2 site contributes to the interaction interface, whereas the Zn1 site stabilises the overall effector fold.

Approximately 10% of all protein structures in the Protein Data Bank contain zinc-binding sites, highlighting the widespread role of zinc in protein stability and function (Ireland and Martin 2019). The most common tetrahedral coordination motifs, CCCC and CCCH, are typically associated with structural stabilisation (Peters et al. 2010; Outram et al. 2022). Although both zinc-binding sites in AvrSr33 adopt canonical tetrahedral geometry (CCCC/CCCH) and contribute to structural stability, our ICP-MS analysis detected approximately one zinc ion per protomer in both AvrSr33-01 and AvrSr33-05, suggesting partial occupancy of zinc-binding sites. *In planta* mutational analyses further indicate that these sites make distinct contributions. Similar functional differentiation of zinc-binding sites has been observed in other zinc-binding effectors. In AvrP, although mutations at the three zinc-binding sites abolish recognition by the immunoreceptor P, Zn1 and Zn2 mutants show reduced accumulation compared with wild-type and Zn3 variants, indicating that different zinc-binding sites contribute unequally to overall structural integrity (Zhang et al. 2018). In AvrSr27, four potential zinc-binding sites are distributed across the N- and C-terminal domains, yet ICP–MS analysis detected only two zinc ions per molecule, indicating incomplete occupancy (Outram et al. 2024). The N-terminal domain, which contains two zinc sites, was sufficient for Sr27 recognition, implying that not all zinc-binding sites contribute equally to stability of the effector or receptor interaction (Outram et al. 2024). These findings suggest that individual zinc-binding sites can make distinct contributions to protein structure, with roles in both maintaining overall protein stability and shaping effector regions recognised by plant immune receptors. In this context, these observations are consistent with the possibility that zinc binding in these effectors is heterogeneous, with binding sites differing in their relative affinity or occupancy. Such heterogeneous zinc occupancy has been reported in other metal-binding proteins and may reflect the presence of multiple binding states that influence protein conformation and function (Grossoehme and Giedroc 2009; He et al. 2020).

Reciprocal substitution of polymorphic residues between AvrSr33-01 and the non-recognised AvrSr33-05 indicates the Zn2-proximal region plays a key role in Sr33 recognition. Mutations within this region abolished recognition of AvrSr33-01 by Sr33, while the corresponding substitutions in AvrSr33-05 restored recognition, supporting a central role for this region in determining recognition specificity. Split GAL4-Ruby protein-protein interaction assays further confirmed that the Zn2-proximal region mediates direct interaction with Sr33. Within this region, the SL/YM polymorphism proved critical for recognition specificity. Substituting SL with YM in AvrSr33-01 abolished Sr33-mediated cell death, whereas the reciprocal change in AvrSr33-05 (YM/SL) reinstated recognition. Consistent with this, sequence alignment of eight AvrSr33 variants revealed strong evolutionary conservation at these positions: all recognized alleles encode SL, while all unrecognized alleles encode YM.

From a structure perspective, these residues are located on a short Zn2-coordinated loop (CSTLS^70^RL^72^PC) defined by two cysteine residues, which position the SL residues on the solvent-exposed surface. In contrast to secondary structure elements such as α-helices and β-sheets, loop regions are inherently flexible, and Zn coordination acts to rigidify their geometry, thereby enabling the formation of a conformation competent for interaction with a plant receptor. The geometry of this loop is restricted by Zn coordination, such that substitutions at the two cysteine residues may perturb local structural integrity. As a result, only a limited number of surface-exposed residues within this loop appear able to tolerate polymorphism, suggesting that Zn coordination imposes strong structural constraints on effector evolution at this recognition interface. Notably, the short helix motif STL^67-69^ within this loop is strictly conserved across all eight variants, whereas substitutions at adjacent positions (R/K^71^and P/A^73^) are tolerated and do not alter recognition, suggesting that sequence variation at this interface is differentially constrained. Protein-protein interaction assays further indicated that the YM-SL substitution in AvrSr33-05 restored recognition, although the weaker betalain pigmentation compared with wild type suggests that additional residues may contribute to interface stability *in planta*. Differences in polarity and side-chain bulk between the YM and SL residues may also subtly influence the conformation of the Zn2-coordinated loop or its interaction with Sr33. Interestingly, a similar pattern is observed in the stem rust effector AvrSr27, where two key residues (K50 and T58) within the Zn2-coordinated loop (K^50^CDSCQLHT^58^) (Fig. S7) are required for Sr27 recognition (Outram et al. 2024). Thus, recognition by Sr27 and Sr33 may depend on the structural conservation of zinc-coordinated loop.

Biochemical and structure analyses indicate that zinc coordination in AvrSr33 primarily contributes to structural integrity. However, these findings do not directly reveal the virulence function of the effector. Notably, the function of pathogen effectors is not to be recognised by host immune receptors, but to promote pathogen fitness within the host. In several bacterial pathogens, zinc-dependent T3SS effectors (Sun et al. 2016; Baruch et al. 2011) share a canonical metalloprotease motif (HEXXH) in which two histidine residues coordinate zinc, which is required for catalytic activity. By contrast, AvrSr33 contains CCCC and CCCH sites, and comparative structural analyses using DALI and Foldseek identified only weak structural similarity between AvrSr33 and diverse β-rich scaffold proteins or domains with no clear catalytic architecture or close structural homolog identified (Fig S8). Together, these observations suggest that zinc coordination in AvrSr33 likely reinforces protein architecture rather than supporting a conserved enzymatic function.

From an evolutionary perspective, our results suggest that Sr33 recognition is focused on a structurally constrained and functionally important region of AvrSr33. More broadly, rather than recognizing arbitrary exposed surface, plant receptors may target structurally conserved and functionally essential regions of effectors that pathogens can least afford to change. For example, the NLR receptor ROQ1 from *N. benthamiana* recognises the *Xanthomonas* effector XopQ by inserting a receptor loop into the effector active-site cleft and targeting conserved residues required for nucleoside-binding, thereby targeting a region of the effector that is constrained by its function (Martin et al. 2020). This evolutionary principle is further reflected in the integrated decoy model, in which plant NLRs have incorporated domains that mimic effector targets. These integrated mimic domains enable receptors to engage the functional sites of effectors, allowing immune recognition of their virulence-associated structural features (Kroj et al. 2016; Sarris et al. 2016; Chen et al. 2022). A classic example is the rice NLR pair RGA5-RGA4, where the integrated heavy metal-associated (HMA) domain of RGA5 mimics the host target of the blast fungus effector Avr-Pia and directly engages its functional interface to trigger RGA4-dependent immunity (Cesari et al. 2014; Ortiz et al. 2017). In our study, this principle is reflected in the recognition of a zinc-anchored loop of AvrSr33-01, where the zinc coordination may constrain the structural framework of this region, while sequence variation within a limited set of exposed surface residues can alter recognition outcomes. The conservation of the STL motif within the Zn2-coordinated loop, together with the retention of the Zn2-binding site across all variants, suggests that this region is subject to constraints beyond simple structure stabilisation. It potentially reflects a functional interface involved in host target interaction, although its precise role remains to be determined.

In summary, four recognised Avr effectors from wheat rust have been shown to coordinate zinc, including AvrSr33, 27, 22 and 13 (Outram et al. 2024; Outram et al. 2026)). Work described here for AvrSr33 provide a structural framework for understanding how zinc coordination governs both effector stability and immune recognition. This zinc-dependent balance between fold integrity and receptor activation exemplifies how plants exploit structural constraints in pathogen effectors to limit evolutionary escape, offering new avenues for rational engineering of disease resistance.

## Acknowledgments

This work was supported by funding from the Australian Research Council (ARC) (Discovery Projects DP240102982). S.J.W. was supported by ARC Future fellowships (FT200100135). D.S.Y. was a recipient of an Australian Government Research Training Programme (RTP) Stipend Scholarship and the Australian Institute of Nuclear Science and Engineering (AINSE) Ltd. Postgraduate Research Award (PGRA). This work was supported by the Commonwealth Scientific and Industrial Research Organization Research Office and the Gatsby Foundation.

The authors acknowledge the use of the ANU crystallisation facility. The authors also acknowledge use of the Australian Synchrotron MX facility and thank the staff for their support. This research was undertaken in part using the MX2 and MX3 beamline at the Australian Synchrotron, part of ANSTO, and made use of the Australian Cancer Research Foundation (ACRF) detector.

## Author contributions

ZL, JC, JS, MF, PND and SJW designed the study; ZL, JC, EMC, MAO, DSY performed the experiments, and all authors analysed the data. ZL, SJW and PND wrote the original manuscript draft, and all authors contributed to writing, reviewing, and editing the manuscript.

## Data availability

The structure of AvrSr33 has been deposited to the protein databank under the PDB code 26OF.

