## Supplemental for "Zinc coordination by the wheat rust effector AvrSr33 provides a structural scaffold required for immune recognition"

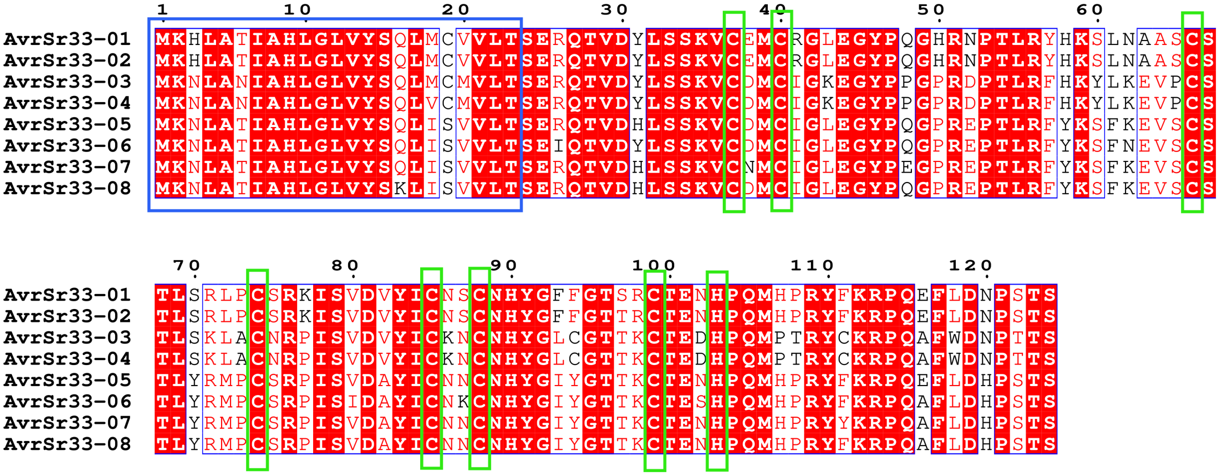

| | AvrSr33-01 | AvrSr33-02 | AvrSr33-03 | AvrSr33-04 | AvrSr33-05 | AvrSr33-06 | AvrSr33-07 | AvrSr33-08 |
|---|---|---|---|---|---|---|---|---|
| AvrSr33-01 | | 99% | 74% | 74% | 81% | 79% | 78% | 79% |
| AvrSr33-02 | 99% | | 75% | 74% | 82% | 80% | 79% | 80% |
| AvrSr33-03 | 74% | 75% | | 99% | 77% | 75% | 77% | 77% |
| AvrSr33-04 | 74% | 74% | 99% | | 77% | 75% | 77% | 77% |
| AvrSr33-05 | 81% | 82% | 77% | 77% | | 94% | 97% | 98% |
| AvrSr33-06 | 79% | 80% | 75% | 75% | 94% | | 93% | 94% |
| AvrSr33-07 | 78% | 79% | 77% | 77% | 97% | 93% | | 97% |
| AvrSr33-08 | 79% | 80% | 77% | 77% | 98% | 94% | 97% | |

**Figure S1:** Alignment and identity sequence matrix of the eight AvrSr33 variants. (a) Blue box indicates the signal peptide, and the green box marks the cysteine and histidine residues forming the zinc-binding sites. The key residues associated with recognition, $SL^{70,72}$/YM are highlighted by black boxes. Recognised and non-recognised variants are separated by dashed lines. (b) Numbers indicate percentage sequence identity between AvrSr33 variants. Darker shading represents higher sequence identity.

**(a) Native sample**

AvrSr33-01
BL21(DE3) pLysS

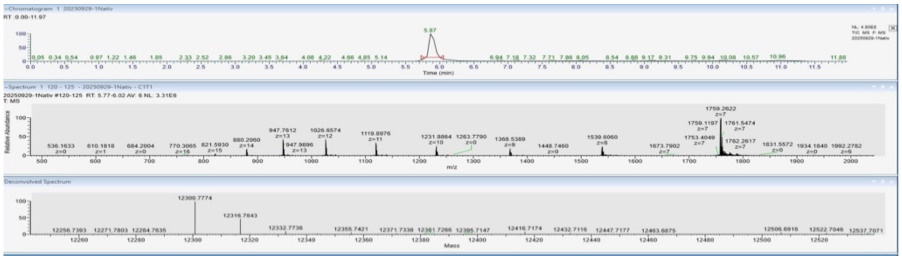

AvrSr33-01
Shuffle

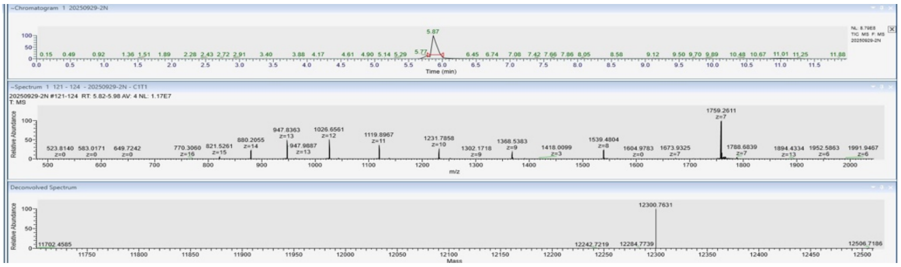

AvrSr33-05
BL21(DE3) pLysS

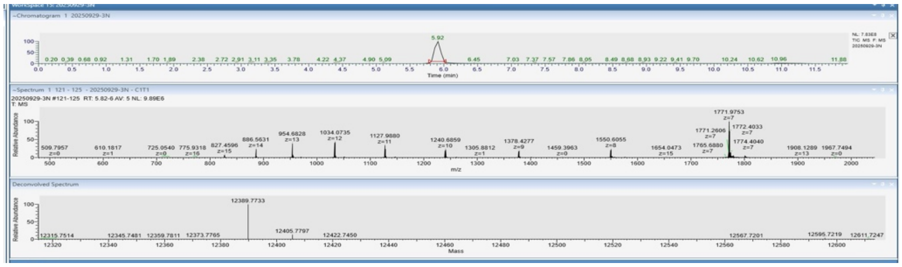

**(b) Reduced sample (treated with DTT)**

AvrSr33-01
BL21(DE3) pLysS

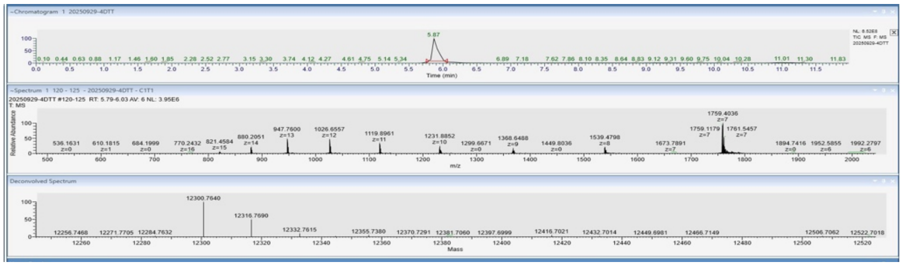

AvrSr33-01
Shuffle

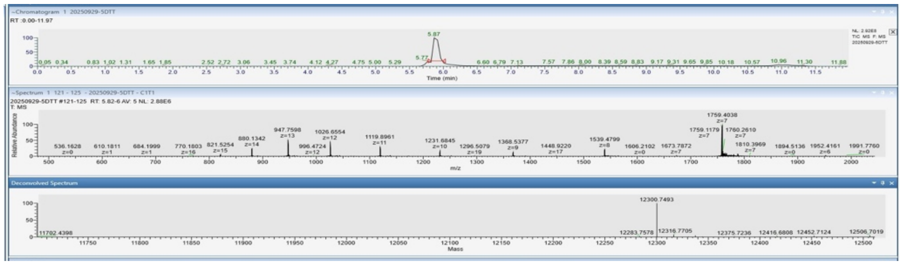

AvrSr33-05
BL21(DE3) pLysS

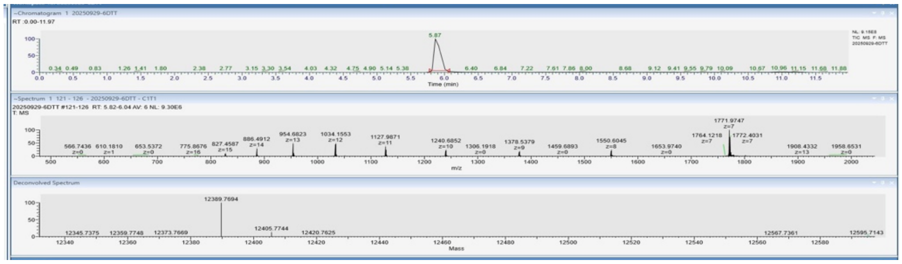

(c) Summary table

| Protein | Theoretical molecular weight (Da) | Native protein | Reduced protein |
|---|---|---|---|
| AvrSr33-01 （BL21 (ED3) pLysS） | 12300.81 | 12300.77 | 12300.76 |
| AvrSr33-01 （Shuffle®） | 12300.81 | 12300.76 | 12300.75 |
| AvrSr33-05 （BL21 (ED3) pLysS） | 12389.08 | 12389.77 | 12389.77 |

**Figure S2:** Intact-mass spectrometry analysis of AvrSr33-01 and AvrSr33-05 proteins expressed in *E.coli* BL21(DE3) pLysS and Shuffle strains. (a) Native proteins (b) Proteins treated with DTT. (c) Summary table comparing mass determined experimentally with theoretical.

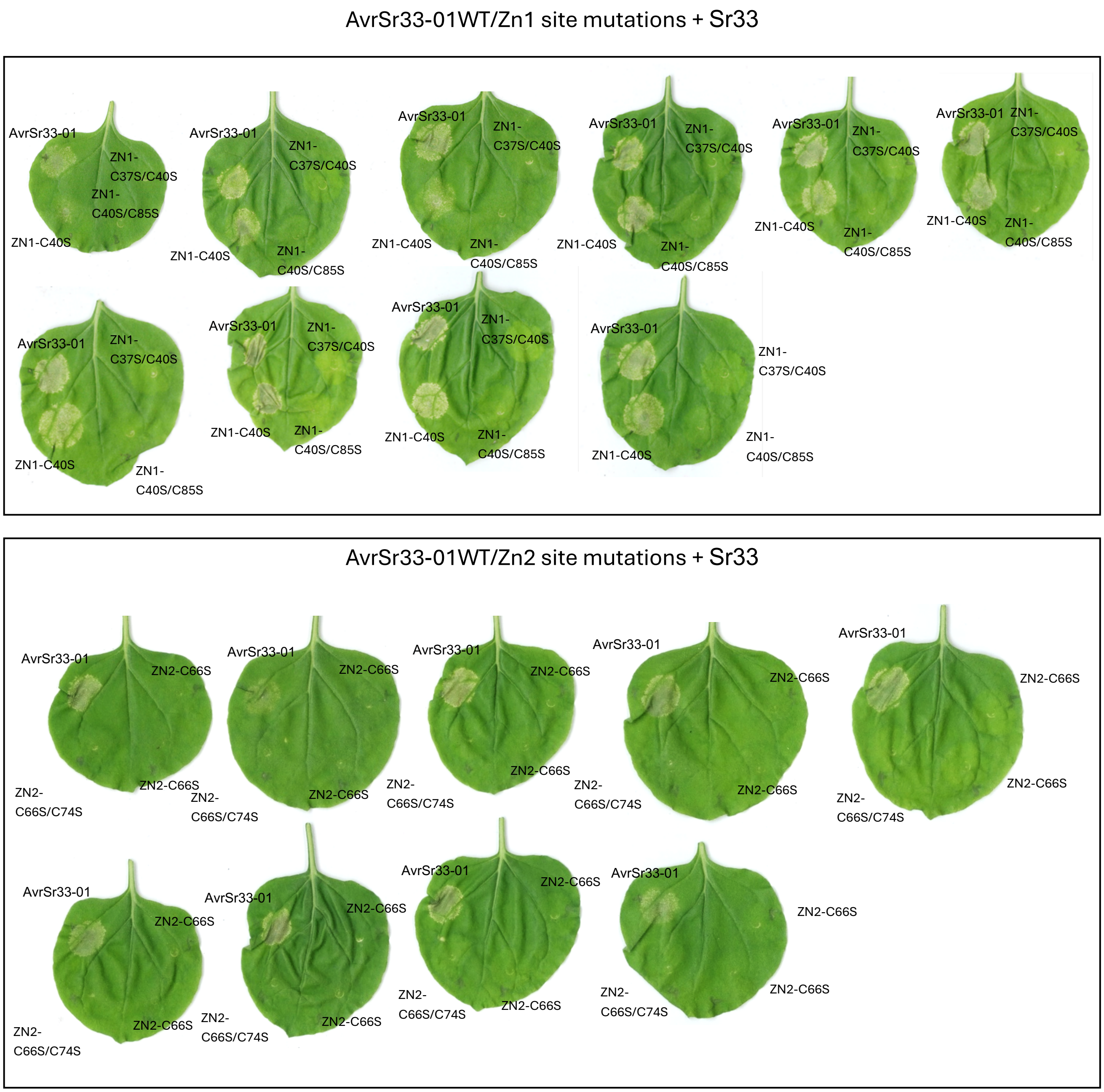

**Figure S3:** Cell death response of *N. benthamiana* leaves co-expressing AvrSr33-01 and its zinc-binding sites mutations with Sr33. (a) AvrSr33-01 and Zn1 site mutations. (b) AvrSr33-01 and Zn2 site mutations.

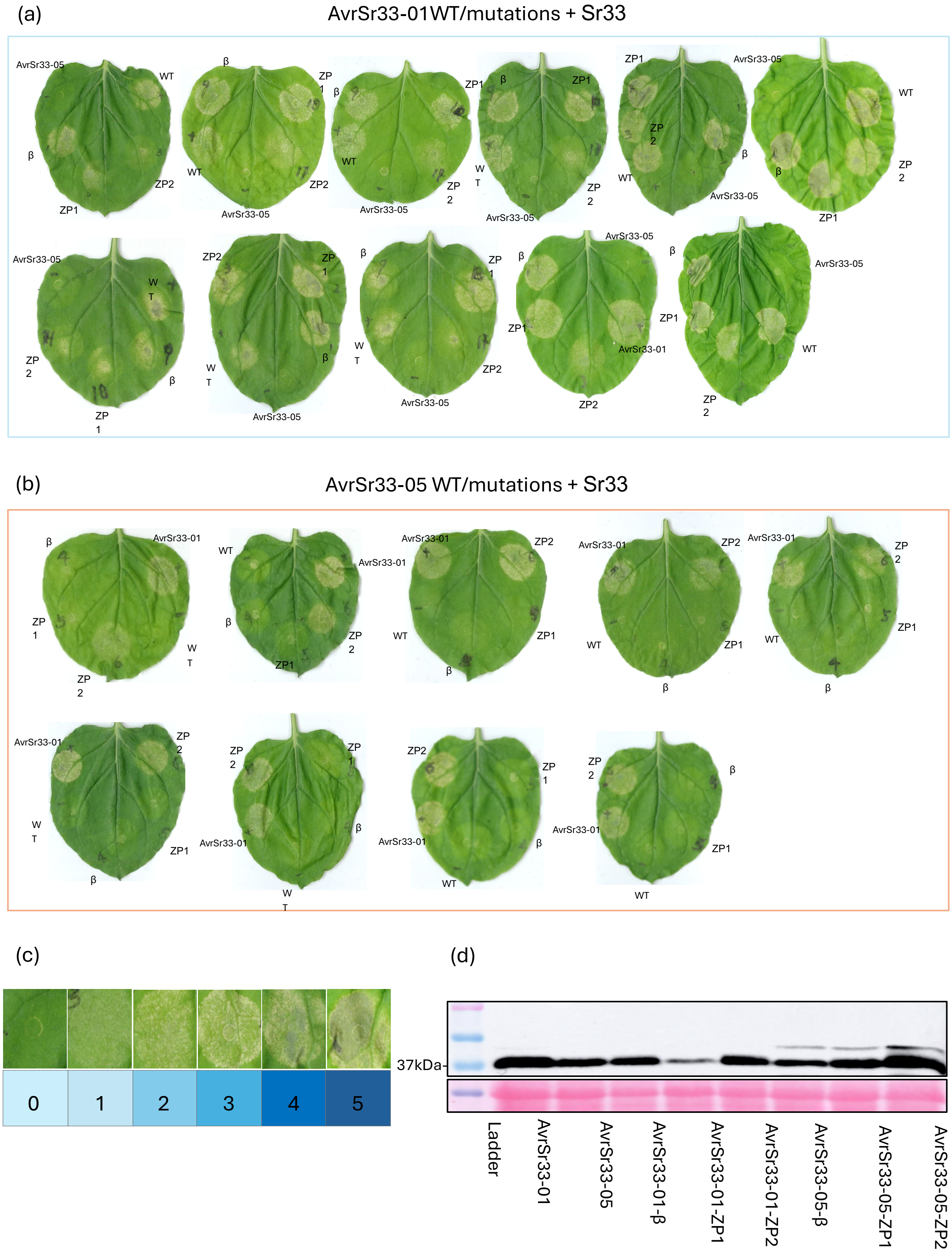

**Figure S4:** Cell death response of *N. benthamiana* leaves co-expressing AvrSr33-01, AvrSr33-05 and their mutations with Sr33. (a) is for AvrSr33-01 and mutations. (b) is for AvrSr33-05 and mutations. (c) Cell death was scored according to the symptoms scale determined from 0-5. AvrSr33-01 and AvrSr33-05 were used as positive and negative control, respectively. (d) Anti-GFP immunoblot of Zn1-proximalinally YFP tagged AvrSr33-01 and AvrSr33-05 and their mutations.

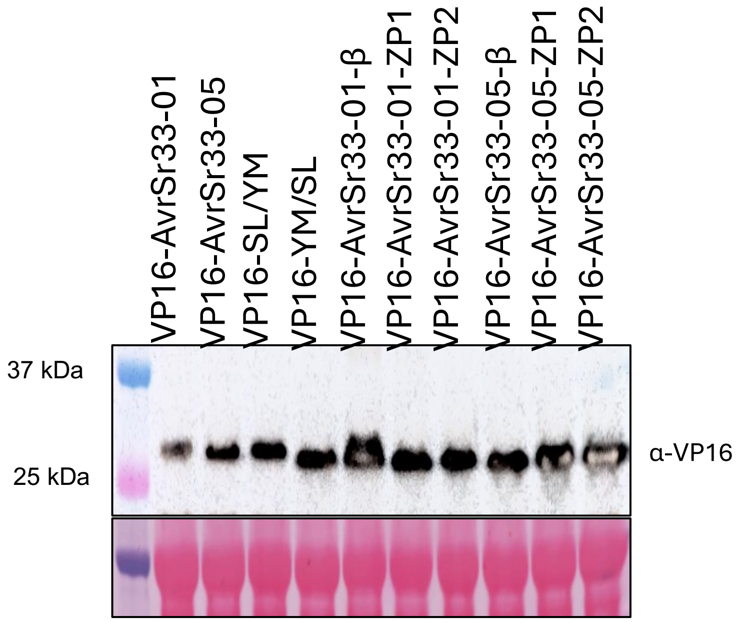

**Figure S5:** Immunoblot analysis confirming accumulation of split-GAL4 RUBY assay. Immunoblots were probed with anti-VP16 antibody. Ponceau S staining Rubisco large subunit (rbcL) was used as a loading control. Comparable accumulation across all constructs confirms consistent protein expression.

(a) Single and multiple amino acid mutations in the Zn2-proximal region of AvrSr33-01 + Sr33

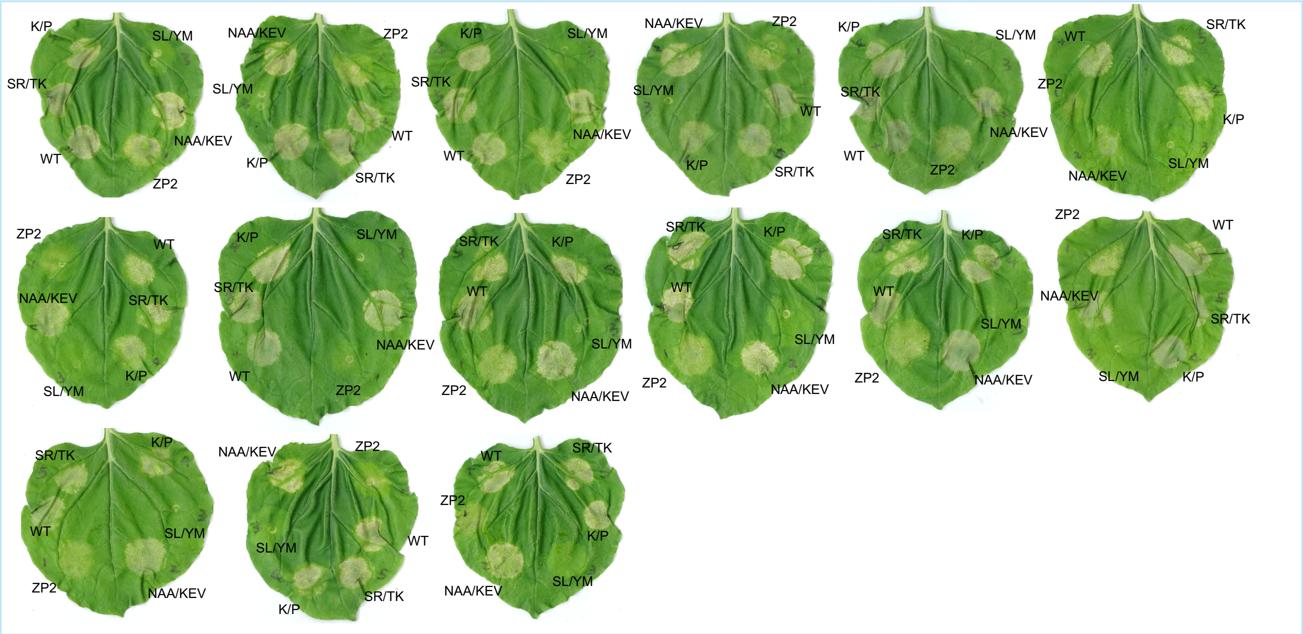

(b) Single and multiple amino acid mutations in the Zn2-proximal region of AvrSr33-05 +Sr33

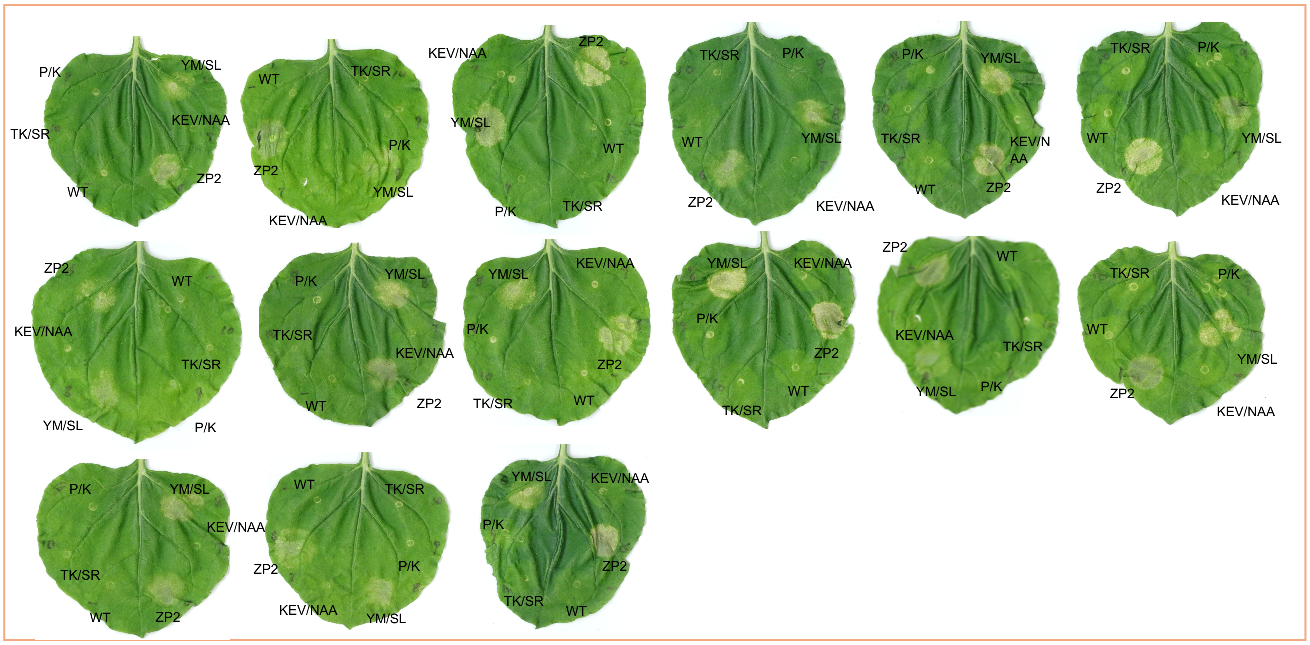

(c)

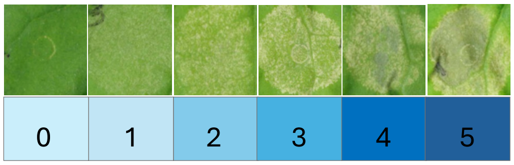

(d)

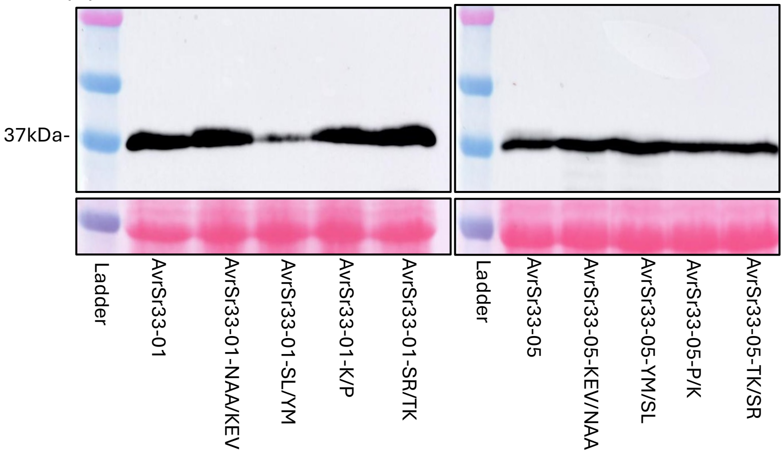

**Figure S6:** Cell death responses of *N. benthamiana* leaves co-expressing AvrSr33-01, AvrSr33-05 and their mutations with Sr33. (a) AvrSr33-01 with single or multiple mutations. (b) AvrSr33-05 with single or multiple mutations. (c) Cell death responses was scored according to the symptoms scale determined from 0-5. AvrSr33-01 and AvrSr33-05 were used as positive and negative control, respectively. (d) Anti-GFP immunoblot of Zn1-proximalinally YFP tagged AvrSr33-01 and AvrSr33-05 and their mutations. Ponceau S staining was used as a loading control.

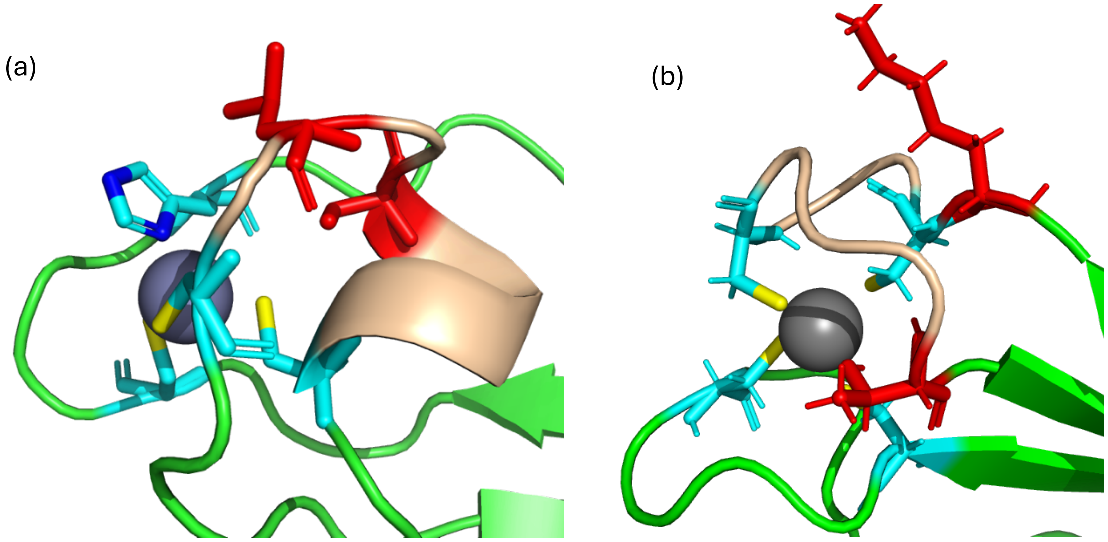

**Figure S7: AvrSr33 structure highlighting the zinc binding sites.** (a) Zn2-loop (CSTLS$^{70}$RL$^{72}$PC) of AvrSr33-01 (b) Zn2-loop (K$^{50}$CDSCQLHT$^{58}$) of AvrSr27. Cysteine residues are shown in cyan, recognition-critical residues in blue, and the remaining residues within the loop are in wheat.

(a)

| PDB | Z-score | RMSD | Aligned length | Seq identity (%) | Description |
|---|---|---|---|---|---|
| 7r78A | 4 | 3.1 | 63 | 11 | molecule: DNA repair protein RAD8; |
| 3ljuX | 3.9 | 5 | 56 | 9 | molecule: ARF-GAP with dual PH domain-containing protein 1; |
| 7tomA | 3.6 | 3.1 | 47 | 11 | molecule: Glycerol dibiphytanyl glycerol tetraether - macro |
| 6i1mA | 3.3 | 3.9 | 65 | 2 | molecule: Cystatin; |
| 8aim-B | 2.8 | 3.3 | 55 | 4 | Uracil-DNA Glycosylase-related inhibitor |

(b)

| Target ID | Description | Aligned length | TM-score | Qcov | RMSD (Å) | E-value | Seq. ID (%) on aligned residues |
|---|---|---|---|---|---|---|---|
| 3tvv-assembly1_A | tandem PH domains of Rtt106 | 54 | 0.409 | 0.48 | 4.59 | 1.7 | 12.7 |
| 2zhx-assembly2_D | Uracil-DNA Glycosylase-related inhibitor | 68 | 0.4688 | 0.61 | 6.5 | 4.77 | 14.2 |
| 1xke-assembly1_A | second Ran-binding domain | 57 | 0.455 | 0.5089286 | 6.92 | 4.77 | 3.2 |
| 1ww4-assembly1_A | cylindracea galectin | 53 | 0.433 | 0.4732143 | 5.56 | 5.38 | 15 |

**Figure S8. Representative structural similarities identified for AvrSr33-01 using DALI and Foldseek searches.** (a) representative structural matches identified by DALI against the PDB database. (b) representative structural matches identified by Foldseek against the PDB100 database. Reported parameters include Z-score, RMSD, aligned length and sequence identity for DALI results, and TM-score, query coverage (Qcov), RMSD, E-value and sequence identity for Foldseek results.

| Name | sequence |
| --- | --- |
| AvrSr 33-01 | GPMVLTSERQTVDYLSSKVCEMCRGLEGYPQGHRNPTLRYHKSLNAASCSTLSRLPCSRKISVDVYICNSCNHYGF FGTSRCTENHPQMHPRYFKRPQEFLDNPSTSS |
| AvrSr 33-05 | GPMVLTSERQTVDHLSSKVCDMCIGLEGYPQGPREPTLRFYKSFKEVSCSTLYRMPCSRPISVDAYICNNCNHYGI YGTTKCTENHPQMHPRYFKRPQEFLDHPSTSS |

**Table S1:** Protein sequences following purification and tag-cleavage.

| Name | Origin | Sequence （5'→3'） |
|---|---|---|
| **AvrSr33-01** | | ATGTCCGAGAGGCAAACCGTGGACTACCTCTCCAGCAAGGTGTGCGAGATGTGCAGGGGACTC<br>GAGGGCTACCCACAAGGCCATAGGAACCCAACTCTCCGCTACCACAAGTCCCTCAACGCCGCCT<br>CTTGCAGCACTCTCTCTAGGCTTCCATGCAGCCGCAAGATCTCCGTGGATGTGTACATCTGCAA<br>CAGCTGCAACCACTACGGATTCTTCGGCACCTCCAGGTGCACCGAGAACCATCCACAAATGCAC<br>CCACGCTACTTCAAGAGGCCACAAGAGTTCCTCGACAACCCATCCACCAGCTGA |
| **AvrSr33_01_Zn1-C40S** | **gBlock** | ATGTCCGAGAGGCAAACCGTGGACTACCTCTCCAGCAAGGTGTGCGAGATGtctAGGGGACTCGAGG<br>GCTACCCACAAGGCCATAGGAACCCAACTCTCCGCTACCACAAGTCCCTCAACGCCGCCTCTTGCAGC<br>ACTCTCTCTAGGCTTCCATGCAGCCGCAAGATCTCCGTGGATGTGTACATCTGCAACAGCTGCAACCA<br>CTACGGATTCTTCGGCACCTCCAGGTGCACCGAGAACCATCCACAAATGCACCCACGCTACTTCAAGA<br>GGCCACAAGAGTTCCTCGACAACCCATCCACCAGCTGA |
| **AvrSr33_01_Zn1-C37/40S** | **gBlock** | ATGTCCGAGAGGCAAACCGTGGACTACCTCTCCAGCAAGGTGTCTGAGATGtctAGGGGACTCGAGG<br>GCTACCCACAAGGCCATAGGAACCCAACTCTCCGCTACCACAAGTCCCTCAACGCCGCCTCTTGCAGC<br>ACTCTCTCTAGGCTTCCATGCAGCCGCAAGATCTCCGTGGATGTGTACATCTGCAACAGCTGCAACCA<br>CTACGGATTCTTCGGCACCTCCAGGTGCACCGAGAACCATCCACAAATGCACCCACGCTACTTCAAGA<br>GGCCACAAGAGTTCCTCGACAACCCATCCACCAGCTGA |
| **AvrSr33_01_Zn1-C40/85S** | **gBlock** | ATGTCCGAGAGGCAAACCGTGGACTACCTCTCCAGCAAGGTGTGCGAGATGtctAGGGGACTCGAGG<br>GCTACCCACAAGGCCATAGGAACCCAACTCTCCGCTACCACAAGTCCCTCAACGCCGCCTCTTGCAGC<br>ACTCTCTCTAGGCTTCCATGCAGCCGCAAGATCTCCGTGGATGTGTACATCtctAACAGCTGCAACCAC<br>TACGGATTCTTCGGCACCTCCAGGTGCACCGAGAACCATCCACAAATGCACCCACGCTACTTCAAGAG<br>GCCACAAGAGTTCCTCGACAACCCATCCACCAGCTGA |
| **AvrSr33_01_Zn2-C66S** | **gBlock** | ATGTCCGAGAGGCAAACCGTGGACTACCTCTCCAGCAAGGTGTGCGAGATGTGCAGGGGACTCGAG<br>GGCTACCCACAAGGCCATAGGAACCCAACTCTCCGCTACCACAAGTCCCTCAACGCCGCCTCTtccAGC<br>ACTCTCTCTAGGCTTCCATGCAGCCGCAAGATCTCCGTGGATGTGTACATCTGCAACAGCTGCAACCA<br>CTACGGATTCTTCGGCACCTCCAGGTGCACCGAGAACCATCCACAAATGCACCCACGCTACTTCAAGA<br>GGCCACAAGAGTTCCTCGACAACCCATCCACCAGCTGA |
| **AvrSr33_01_Zn2-C66/74S** | **gBlock** | ATGTCCGAGAGGCAAACCGTGGACTACCTCTCCAGCAAGGTGTGCGAGATGTGCAGGGGACTCGAG<br>GGCTACCCACAAGGCCATAGGAACCCAACTCTCCGCTACCACAAGTCCCTCAACGCCGCCTCTtccAGC<br>ACTCTCTCTAGGCTTCCAtctAGCCGCAAGATCTCCGTGGATGTGTACATCTGCAACAGCTGCAACCAC<br>TACGGATTCTTCGGCACCTCCAGGTGCACCGAGAACCATCCACAAATGCACCCACGCTACTTCAAGAG<br>GCCACAAGAGTTCCTCGACAACCCATCCACCAGCTGA |
| **AvrSr33-01_beta sheet** | **gBlock** | ATGTCCGAGAGGCAAACCGTGGACCACCTCTCCAGCAAGGTGTGCGAGATGTGCAGGGGACTCGAG<br>GGCTACCCACAAGGCCATAGGAACCCAACTCTCCGCTTCTACAAGTCCTTCAACGCCGCCTCTTGCAG<br>CACTCTCTCTAGGCTTCCATGCAGCCGCAAGATCTCCGTGGATGCCTACATCTGCAACAGCTGCAACC<br>ACTACGGAATCTACGGCACCTCCAGGTGCACCGAGAACCATCCACAAATGCACCCACGCTACTTCAA<br>GAGGCCACAAGAGTTCCTCGACCATCCATCCACCAGCTGA |
| **AvrSr33-01_ZP1** | **gBlock** | ATGTCCGAGAGGCAAACCGTGGACTACCTCTCCAGCAAGGTGTGCGATATGTGCATCGGACTCGAG<br>GGCTACCCACAAGGCCCAAGGGAGCCAACTCTCCGCTACCACAAGTCCCTCAACGCCGCCTCTTGCA<br>GCACTCTCTCTAGGCTTCCATGCAGCCGCAAGATCTCCGTGGATGTGTACATCTGCAACAACTGCAAC<br>CACTACGGATTCTTCGGCACCTCCAGGTGCACCGAGAACCATCCACAAATGCACCCACGCTACTTCAA<br>GAGGCCACAAGAGTTCCTCGACAACCCATCCACCAGCTGA |
| **AvrSr33-01_ZP2** | **gBlock** | ATGTCCGAGAGGCAAACCGTGGACTACCTCTCCAGCAAGGTGTGCGAGATGTGCAGGGGACTCGAG<br>GGCTACCCACAAGGCCATAGGAACCCAACTCTCCGCTACCACAAGTCCCTCAAGGAAGTGTCTTGCA<br>GCACTCTCTACAGGATGCCATGCAGCCGCCCAATCTCCGTGGATGTGTACATCTGCAACAGCTGCAAC<br>CACTACGGATTCTTCGGCACCACCAAGTGCACCGAGAACCATCCACAAATGCACCCACGCTACTTCAA<br>GAGGCCACAAGAGTTCCTCGACAACCCATCCACCAGCTGAg |
| **AvrSr33-01_K$^{77}$/P** | **gBlock** | ATGTCCGAGAGGCAAACCGTGGACTACCTCTCCAGCAAGGTGTGCGAGATGTGCAGGGGACTCGAG<br>GGCTACCCACAAGGCCATAGGAACCCAACTCTCCGCTACCACAAGTCCCTCAACGCCGCCTCTTGCAG<br>CACTCTCTCTAGGCTTCCATGCAGCCGCccaATCTCCGTGGATGTGTACATCTGCAACAGCTGCAACCA<br>CTACGGATTCTTCGGCACCTCCAGGTGCACCGAGAACCATCCACAAATGCACCCACGCTACTTCAAGA<br>GGCCACAAGAGTTCCTCGACAACCCATCCACCAGCTGA |

| | | |
|---|---|---|
| **AvrSr33-01_SR$^{97,98}$/TK** | **gBlock** | ATGTCCGAGAGGCAAACCGTGGACTACCTCTCCAGCAAGGTGTGCGAGATGTGCAGGGGACTCGAGGGCTACCCACAAGGCCATAGGAACCCAACTCTCCGCTACCACAAGTCCCTCAACGCCGCCTCTTGCAGCACTCTCTCTAGGCTTCCATGCAGCCGCAAGATCTCCGTGGATGTGTACATCTGCAACAGCTGCAACCACTACGGATTCTTCGGCACCaccaagTGCACCGAGAACCATCCACAAATGCACCCACGCTACTTCAAGAGGCCACAAGAGTTCCTCGACAACCCATCCACCAGCTGA |
| **AvrSr33-05** | | ATGTCCGAGAGGCAAACCGTGGACCACCTCTCCAGCAAGGTGTGCGATATGTGCATCGGCCTCGAGGGCTACCCACAAGGCCCAAGAGAGCCAACTCTCAGGTTCTACAAGTCCTTCAAGGAAGTGTCCTGCAGCACCCTCTACAGGATGCCATGCTCCAGGCCAATCTCCGTGGACGCCTACATCTGCAACAACTGCAACCACTACGGCATCTACGGCACCACCAAGTGCACCGAGAACCATCCACAAATGCACCCACGCTACTTCAAGAGGCCACAAGAGTTCCTCGACCATCCATCCACCAGC |
| **AvrSr33-05_beta sheet** | **gBlock** | ATGTCCGAGAGGCAAACCGTGGACTACCTCTCCAGCAAGGTGTGCGATATGTGCATCGGCCTCGAGGGCTACCCACAAGGCCCAAGAGAGCCAACTCTCAGGTACCACAAGTCCCTCAAGGAAGTGTCCTGCAGCACCCTCTACAGGATGCCATGCTCCAGGCCAATCTCCGTGGACGTGTACATCTGCAACAACTGCAACCACTACGGCTTCTTCGGCACCACCAAGTGCACCGAGAACCATCCACAAATGCACCCACGCTACTTCAAGAGGCCACAAGAGTTCCTCGACAACCCATCCACCAGCTGA |
| **AvrSr33-05_ZP1** | **gBlock** | ATGTCCGAGAGGCAAACCGTGGACCACCTCTCCAGCAAGGTGTGCGAGATGTGCAGGGGCCTCGAGGGCTACCCACAAGGCCATAGAAACCCAACTCTCAGGTTCTACAAGTCCTTCAAGGAAGTGTCCTGCAGCACCCTCTACAGGATGCCATGCTCCAGGCCAATCTCCGTGGACGCCTACATCTGCAACAGCTGCAACCACTACGGCATCTACGGCACCACCAAGTGCACCGAGAACCATCCACAAATGCACCCACGCTACTTCAAGAGGCCACAAGAGTTCCTCGACCATCCATCCACCAGCTGA |
| **AvrSr33-05_ZP2** | **gBlock** | ATGTCCGAGAGGCAAACCGTGGACCACCTCTCCAGCAAGGTGTGCGATATGTGCATCGGCCTCGAGGGCTACCCACAAGGCCCAAGAGAGCCAACTCTCAGGTTCTACAAGTCCTTCAACGCCGCCTCCTGCAGCACCCTCTCTAGGCTTCCATGCTCCAGGAAGATCTCCGTGGACGCCTACATCTGCAACAACTGCAACCACTACGGCATCTACGGCACCTCCAGGTGCACCGAGAACCATCCACAAATGCACCCACGCTACTTCAAGAGGCCACAAGAGTTCCTCGACCATCCATCCACCAGCTGA |
| **AvrSr33-05_P$^{77}$/K** | **gBlock** | ATGTCCGAGAGGCAAACCGTGGACCACCTCTCCAGCAAGGTGTGCGATATGTGCATCGGCCTCGAGGGCTACCCACAAGGCCCAAGAGAGCCAACTCTCAGGTTCTACAAGTCCTTCAAGGAAGTGTCCTGCAGCACCCTCTACAGGATGCCATGCTCCAGGaagATCTCCGTGGACGCCTACATCTGCAACAACTGCAACCACTACGGCATCTACGGCACCACCAAGTGCACCGAGAACCATCCACAAATGCACCCACGCTACTTCAAGAGGCCACAAGAGTTCCTCGACCATCCATCCACCAGCTGA |
| **AvrSr33-05_TK$^{97,98}$/SR** | **gBlock** | ATGTCCGAGAGGCAAACCGTGGACCACCTCTCCAGCAAGGTGTGCGATATGTGCATCGGCCTCGAGGGCTACCCACAAGGCCCAAGAGAGCCAACTCTCAGGTTCTACAAGTCCTTCAAGGAAGTGTCCTGCAGCACCCTCTACAGGATGCCATGCTCCAGGCCAATCTCCGTGGACGCCTACATCTGCAACAACTGCAACCACTACGGCATCTACGGCACCtccaggTGCACCGAGAACCATCCACAAATGCACCCACGCTACTTCAAGAGGCCACAAGAGTTCCTCGACCATCCATCCACCAGCTGA |
| **Primer** | **Template** | **Sequence** |
| **AvrSr33-01_NAA/KEV-F** | **AvrSr33-01** | CGCTACCACAAGTCCCTCAAGGAAGTGTCTTGCAGCACTCTCTCT |
| **AvrSr33-01_NAA/KEV-R** | | AGAGAGAGTGCTGCAAGACACTTCCTTGAGGGACTTGTGGTAGCG |
| **AvrSr33-01_SL/YM-F** | | GCCTCTTGCAGCACTCTCTACAGGATGCCATGCAGCCGCAAGATC |
| **AvrSr33-01_SL/YM-R** | | GATCTTGCGGCTGCATGGCATCCTGTAGAGAGTGCTGCAAGAGGC |
| **AvrSr33-05_KEV/NAA-F** | **AvrSr33-05** | AGGTTCTACAAGTCCTTCAACGCCGCCTCCTGCAGCACCCTCTAC |

| | | |
|---|---|---|
| **AvrSr33-05_KEV/NAA-R** | | GTAGAGGGTGCTGCAGGAGGCGGCGTTGAAGGACTTGTAGAACCT |
| **AvrSr33-05_YM/SL-F** | | AGTGTCCTGCAGCACCCTCTCTAGGCTTCCATGCTCCAGGCCAATC |
| **AvrSr33-05_YM/SL-R** | | GATTGGCCTGGAGCATGGAAGCCTAGAGAGGGTGCTGCAGGACACT |

**Table S2:** Constructs and primer sequences used in this study.

| Aimless | | | |
|---|---|---|---|
| Unit cell lengths (Å) | 33.85, 39.63, 76.79 | | |
| Unit cell angles (°) | 90, 90, 90 | | |
| Spacegroup | P 21 21 21 | | |
| Wavelength (Å) | 0.95372 | | |
| Wilson B-factor ($A^{-2}$) | 29.83 | | |
| Low resolutiom limit (Å) | 39.63 | | |
| High resolutiom limit (Å) | 1.85 | | |
| | **Overall** | **Innershell** | **Outershell** |
| Total number of observations | 116239 | 920 | 7233 |
| Total number unique | 9343 | 109 | 570 |
| Multiplicity | 12.4 | 8.4 | 12.7 |
| Completeness (%) | 100 | 98.3 | 100 |
| Mean ((I)/sd(I)) | 19.4 | 47.1 | 2.60 |
| Rmerge (all I+ & I-) | 0.065 | 0.035 | 0.475 |
| Rmeas (all I+ & I-) | 0.068 | 0.037 | 0.496 |
| Rpim (all I+ & I-) | 0.190 | 0.012 | 0.140 |
| CC (1/2) | 0.995 | 0.999 | 0.972 |
| **Phenix Refine** | | | |
| Reflections used overall | 9289 | | |
| Reflections used high resolution shell | 3030 | | |
| Reflections used for R-free overall | 469 | | |
| Reflections used for R-free high resolution shell | 151 | | |
| R-work overall | 0.23 | | |
| R-work high resolution shell | 0.26 | | |
| R-free overall | 0.27 | | |
| R-free high resolution shell | 0.33 | | |
| Number of non-hydrogen atoms | 775 | | |
| macromolecules | 749 | | |
| solvent | 24 | | |
| Protein residues | 93 | | |
| Average B-factor ($A^{-2}$) | 40.07 | | |
| macromolecules | 40.08 | | |
| ligands | 30.48 | | |
| solvent | 40.69 | | |
| **MolProbity Statistics** | | | |
| Ramachandran outliers (%) | 0 | | |
| Ramachandran favoured (%) | 97.75 | | |
| Rotamer outliers (%) | 0 | | |
| C-beta deviations | 0 | | |
| Clashscore | 2.04 | | |
| RMS (bonds) | 0.007 | | |
| RMS (angles) | 0.89 | | |
| MolProbity overall score | 1.03 | | |

**Table S3:** X-ray data collection, structure solution and refinement statistics for AvrSr33
